# Copula modeling of gene coexpression in single-cell RNA sequencing data

**DOI:** 10.64898/2025.12.04.692380

**Authors:** Connor Puritz, Rosemary Braun

## Abstract

Single-cell RNA sequencing (scRNA-seq) has become an indispensable tool for studying biological systems at the cellular level. It has therefore has become increasingly important to develop accurate statistical models of scRNA-seq data. While many models have been proposed to characterize transcript expression of individual genes, comparatively little attention has been paid to modeling gene coexpression. Copula modeling offers a flexible approach to modeling gene coexpression by linking models of individual genes together using copula functions. Despite the growing popularity of copula models, their utility for modeling scRNA-seq data has not been thoroughly explored. Here we evaluated six copula models on their ability to model gene coexpression in scRNA-seq data. Using a diverse collection of reference datasets, we evaluated each copula model’s accuracy and efficiency in reproducing gene coexpression patterns. Our results show that Gaussian copulas provide the best balance between accuracy and speed, with more flexible but expensive copula models providing only a marginal improvement in accuracy while requiring a much longer time to fit. Vine copulas show promise in being able to achieve high accuracy, but current implementations are unable to scale to the large size of typical scRNA-seq datasets.

## 1 Introduction

Since it was first developed, single-cell RNA sequencing (scRNA-seq) has become a popular tool for analyzing biological systems on the cellular level by allowing quantification of the transcriptome of individual cells. Compared to other technologies for transcriptome analysis, scRNA-seq presents a number of unique challenges. The number of cells profiled is often in the thousands to millions, several orders of magnitude larger than the sample sizes of bulk RNA-seq or microarray experiments. Furthermore, the transcript counts are much sparser, with expression observed for only a small percentage of genes in each cell (Zappia et al., 2017), and the data may also suffer from zero-inflation. This has led to the development of a plethora of methods designed specifically to address these characteristics of scRNA-seq data.

A common starting point for many scRNA-seq methods is the construction of a statistical model of transcript counts in individual cells. It is important to not only model the expression of individual genes but also the coexpression of genes, as analysis of gene coexpression can provide insight to many important biological processes (e.g. Stuart et al., 2003; Ho et al., 2007; Braun et al., 2008; Gaiteri et al., 2014; Ruprecht et al., 2017; Panwar et al., 2021; Wang et al., 2021). However, many of the statistical models of scRNA-seq data that have been proposed either focus on modeling transcript counts for individual genes (e.g. Zappia et al., 2017; Li and Li, 2019) or do not explicitly model coexpression (e.g. Risso et al., 2018). The latter is important if one wishes to perform in-silico perturbations with a ground truth.

One approach for explicitly modeling coexpression is copula modeling. First introduced by Sklar (1959), copulas can be formally defined as multivariate cumulative distribution functions with uniform margins on the unit interval (Genest et al., 2024). Their utility arises from Sklar’s theorem, which states that any multivariate distribution function can be expressed in terms of its marginal distribution functions and a copula function (Sklar, 1959; Genest et al., 2024). Copulas thus link one-dimensional distributions together to yield multivariate distributions (Nelsen, 2006). Copulas provide an approach for modeling and analyzing dependencies in high-dimensional datasets, and can also be used to construct flexible high-dimensional distributions with a specified dependence structure.

In recent years, copulas have become a popular tool in omics research for modeling dependencies between genes. Zhang (2017) implemented a Bayes classifier of bulk RNA-seq data using a Gaussian copula model of transcript counts. Zhang and Shi (2017) constructed a Gaussian mixture copula model for inferring a Bayesian network from multimodal genomics data. Ray et al. (2020) proposed a copula-based statistical test for detecting differential coexpression of genes from bulk RNA-seq data. Sarkar et al. (2024) proposed a method for inferring cell-cell interactions from spatial transcriptomics data using a Gaussian copula to model joint ligand-receptor expression. ESCO (Tian et al., 2021), SPsimSeq (Assefa et al., 2020), and scDesign2 (Sun et al., 2021) are all methods for generating synthetic scRNA-seq data which use a Gaussian copula model. scDesign3 (Song et al., 2024), which extends scDesign2, generates synthetic single-cell and spatial omics data using both Gaussian and vine copula models.

Despite the extensive use of copulas in omics methods, no systematic evaluation has been performed to assess different copula models on their utility and accuracy in modeling gene dependencies. Instead, the choice of copula model has often been guided by convenience or simplicity. Here, we sought to fill this gap by evaluating six copula models on their performance in modeling gene coexpression patterns in scRNA-seq data. We fit copula models to 38 scRNA-seq datasets covering a range of cell types, tissues, organisms, and sequencing protocols. We then compared models based on their accuracy on several tasks, as well as their scalability. Finally, we give users recommendations for choosing copula models based on the priorities of their analyses.

## 2 Materials and Methods

### 2.1 Copula modeling of scRNA-seq data

Suppose that we have a scRNA-seq dataset with transcript counts for *m* genes from *n* cells. Let **X** = (*X*_1_, …, *X*_*m*_) be a random variable representing the unnormalized transcript counts across all *m* genes for a single cell. The support of each *X*_*i*_ is the nonnegative integers. If there are no covariates to model (e.g. multiple cells types, multiple donors, etc.), we can equate the *m × n* count matrix with a set of *n* independent and identically distributed (iid) samples of **X**.^1^ Given a parametric family of distribution functions {*H*_*θ*_ | *θ* ∈ Θ}, our goal is to learn from the dataset the parameter *θ* such that *H*_*θ*_ best approximates the true distribution function *H* of **X**.

According to Sklar’s theorem (Sklar, 1959), there exists a function *C*: [0, 1]^*m*^ → [0, 1] called a copula such that the distribution function of **X** can be written as

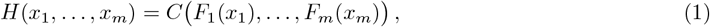

where *F*_*i*_ is the distribution function of the *i*th margin (i.e. the transcript counts for a single gene). Learning *H* is thus equivalent to learning the marginal distributions and the associated copula. Numerous parametric models for transcript counts of individual genes in scRNA-seq datasets have been proposed (e.g. Zappia et al., 2017; Risso et al., 2018). However, as our goal is to evaluate the copula estimators, we chose to model the marginal distributions empirically, since misspecification of the marginal models could degrade the quality of the copula estimators (see Section S1).

In the absence of a ground truth model for a given dataset, it is difficult to evaluate if one copula model is better than another by only examining the copula functions. Instead, we reasoned that if a distribution function *H*_*θ*_ is closer to the true distribution function than another distribution function *H*_*θ*_*′*, then samples drawn from *H*_*θ*_ should be harder to statistically distinguish from the reference dataset than samples drawn from *H*_*θ*_*′*. We thus constructed a diverse compendium of scRNA-seq datasets. For each dataset, we fit a variety of copula models, sampled synthetic datasets from each model, and compared the samples with the reference dataset using a variety of metrics (Figure 1). Below, we describe each of these steps in detail.

**Figure 1:**
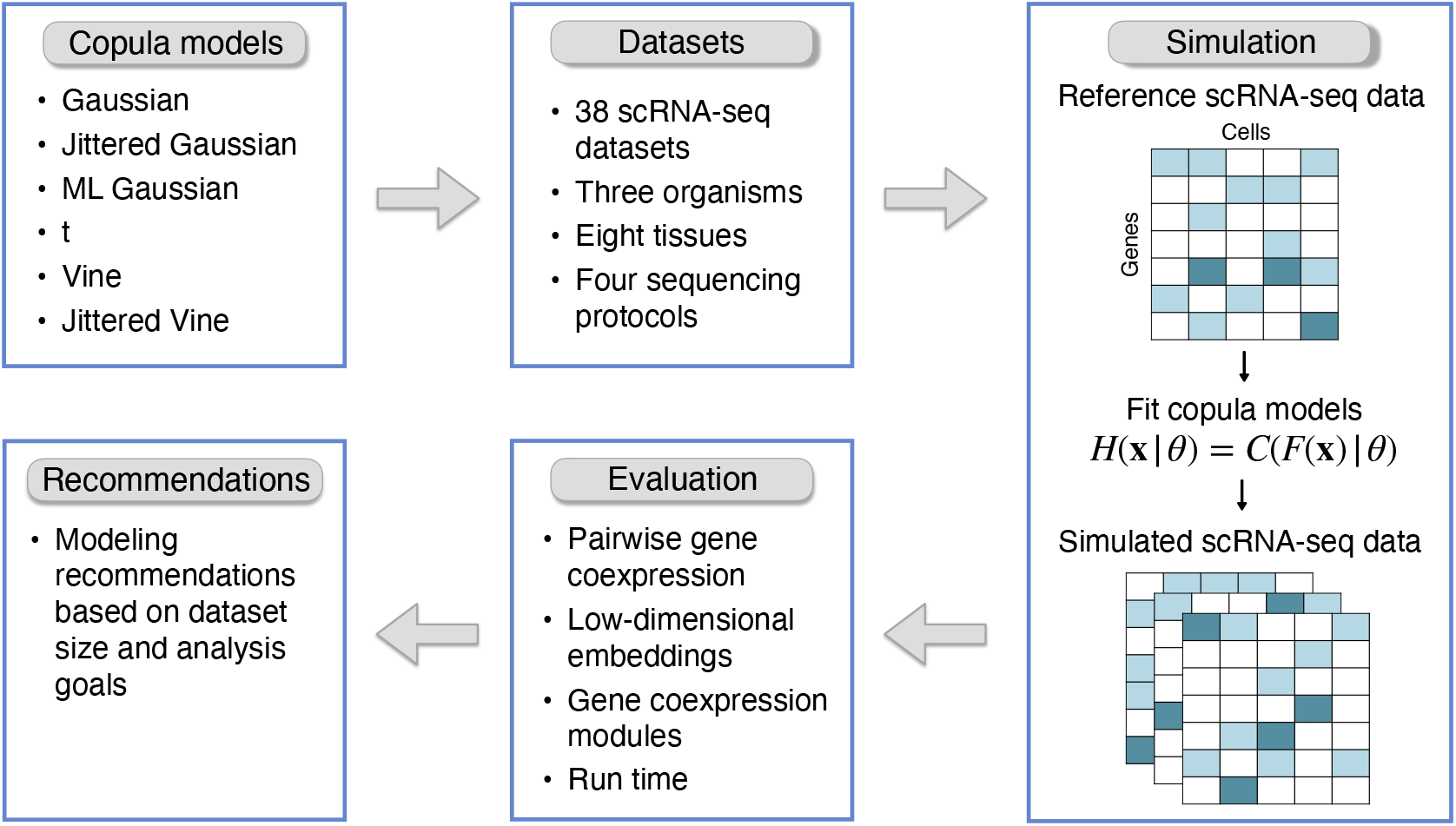
Schematic of the workflow used to evaluate copulas for modeling gene coexpression in single-cell RNA sequencing data.

### 2.2 Copula families

Here we describe the various families of copulas that we evaluated.

#### 2.2.1 Gaussian copula

The Gaussian copula is the copula of the multivariate Gaussian distribution. It is defined as

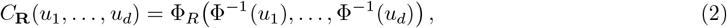

where **R** is a *d × d* correlation matrix, **Φ**^−1^ is the standard normal quantile function, and **Φ**_**R**_ is the distribution function of a *d*-dimensional Gaussian random variable with mean **0** and covariance matrix **R**. The Gaussian copula is one of the most commonly used copulas as it can be easily constructed in arbitrary dimensions and it is intuitively parametrized by a correlation matrix. Gaussian copulas have been previously used for modeling scRNA-seq data (Zhang, 2017; Assefa et al., 2020; Tian et al., 2021; Sun et al., 2021; Song et al., 2024; Sarkar et al., 2024). We consider three different estimators for the correlation matrix: a sample correlation matrix computed using transcript counts (see Section S3.1), a sample correlation matrix computed using jittered margins (see Section 2.2.4, Section S2), and the maximum likelihood estimator of the correlation matrix (see Section S3.2). We refer to these as Gaussian, jittered Gaussian, and ML Gaussian copulas, respectively.

#### 2.2.2 *t* copula

A common criticism of the Gaussian copula is that it does not adequately model tail dependence, i.e., the cooccurrence of extreme events in the tails of marginal distributions (Zeevi and Mashal, 2002). An extension of the Gaussian copula which does model tail dependence is the *t* copula, which is the copula of the multivariate *t* distribution. Similar to the Gaussian copula, the *t* copula is implicitly defined as

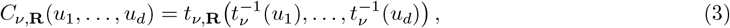

where *ν* is the degrees of freedom; **R** is a correlation matrix; 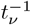 is the quantile function a univariate *t* distribution with degrees of freedom *ν*; and *t*_*ν*,**R**_ is the distribution function of a *d*-dimensional *t* random variable with mean **0**, degrees of freedom *ν*, and scale matrix **R**. Note that the *t* copula converges uniformly to the Gaussian copula in the limit *ν* → ∞ for fixed **R** (Zeevi and Mashal, 2002). To the best of our knowledge, the *t* copula has not previously been applied to omics data. We fit *t* copulas to transcript counts using a two-stage maximum likelihood approach (see Section S3.3).

#### 2.2.3 Vine copulas

Many parametric families of bivariate copulas have been described, each capturing a wide range of dependence structures. While most can be generalized to higher dimensions, these generalizations are often quite inflexible (Czado and Nagler, 2022). Pair-copula decompositions address this by decomposing high-dimensional copulas in terms of bivariate, or pair, copulas. The most popular pair-copula decomposition is the vine copula decomposition, which exploits the fact that any multivariate density function can be decomposed into a product of bivariate conditional density functions. To illustrate this, we consider the following example from Czado and Nagler (2022). Suppose **X** is a three-dimensional random variable (e.g., a trio of coexpressed genes) with joint distribution function *F*, marginal distribution functions *F*_*i*_ (*i* = 1, 2, 3), and copula *C*. Denote the corresponding density functions as *f, f*_*i*_, and *c*, respectively. By Bayes’ theorem, the joint density function can be decomposed as

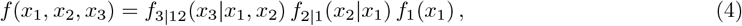

where *f*_*j*|*D*_ denotes the conditional density function of *X*_*j*_ given {*X*_*k*_ = *x*_*k*_ | *k* ∈ *D*} for nonempty *D* ⊆ { 1, 2, 3}. Note that this decomposition is not unique, and different orders of decomposition will give different copulas. We can rewrite Equation 4 in terms of pair copula densities as

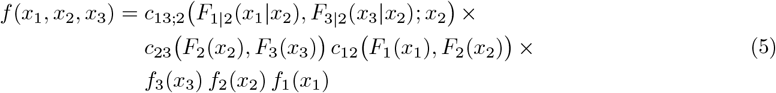

where *c*_13;2_ denotes the copula density function of the conditional distribution of (*X*_1_, *X*_3_) given *X*_2_ = *x*_2_. In practice, the dependence on *x*_2_ in *c*_13;2_ is dropped to facilitate estimation, thus making vine copulas a construction rather than a decomposition (Czado and Nagler, 2022). The construction process involves selecting an order of decomposition, selecting a parametric family for each pair copula, and estimating the parameters for each pair copula. Vine copulas have been previously used to model scRNA-seq data (Fuetterer et al., 2019; Song et al., 2024). We model vine copulas using rvinecopulib (Nagler and Vatter, 2023) and evaluated vine copulas fit directly to transcript count data as well as vine copulas fit using jittered margins (see Section 2.2.4, Section S2, and Section S3.4).

#### 2.2.4 Jittered copulas

There are two possible concerns with copula modeling of discrete data. The first is that the copula of a discontinuous distribution is not identifiable. Sklar’s theorem (Sklar, 1959) states that if the margins of the joint distribution are all continuous, then there exists a unique copula satisfying Equation 1. However, if at least one margin is discontinuous, then there exist infinitely many copulas that satisfy Equation 1 (Nasri and Remillard, 2023). Nevertheless, while the copula may not be identifiable, copula parameters within a fixed copula family often are, and thus parameter estimation is not inherently problematic (see Nasri and Remillard (2023) for a more detailed discussion). The second more practical concern is that parameter estimation is generally more computationally expensive for discrete data than continuous data, and especially so for count data (Panagiotelis et al., 2012). One approach to address these concerns is to “jitter” discrete distributions through the introduction of uniform random noise to make the margins continuous (see Section S2). Sun et al. (2021) used jittered Gaussian copulas for scRNA-seq modeling, while Song et al. (2024) used both jittered Gaussian and jittered vine copulas.

#### 2.2.5 Independence copula

As a baseline for comparison, we also evaluated the independence copula, which is defined as

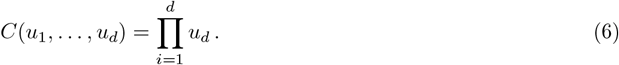

If a distribution’s copula is the independence copula, then the distribution’s margins are independent. When used to model scRNA-seq data, this would imply the expression of each gene has no dependence on the expression of any other gene.

### 2.3 Datasets

Ten publicly-available scRNA-seq studies were selected to construct reference datasets (Table 1). Eight of the studies sequenced human (*Homo sapiens*) cells from a total of seven different tissues. The other two studies sequenced mouse (*Mus musculus*) cells and pig (*Sus scrofa domesticus*) cells. The studies used either a 10x 3’ assay, a 10x 5’ assay, or both. Each study was broken down into several datasets, with each dataset serving as a single reference dataset for simulation. A total of 38 reference datasets were constructed. To minimize covariates that would require more complex modeling, and to ensure that the iid assumptions were met, all datasets were chosen to be from a single donor/sample, of a single cell type, and generated using the same sequencing protocol. In each dataset, genes with zero expression across all cells were removed and counts were log-normalized.

**Table 1:**
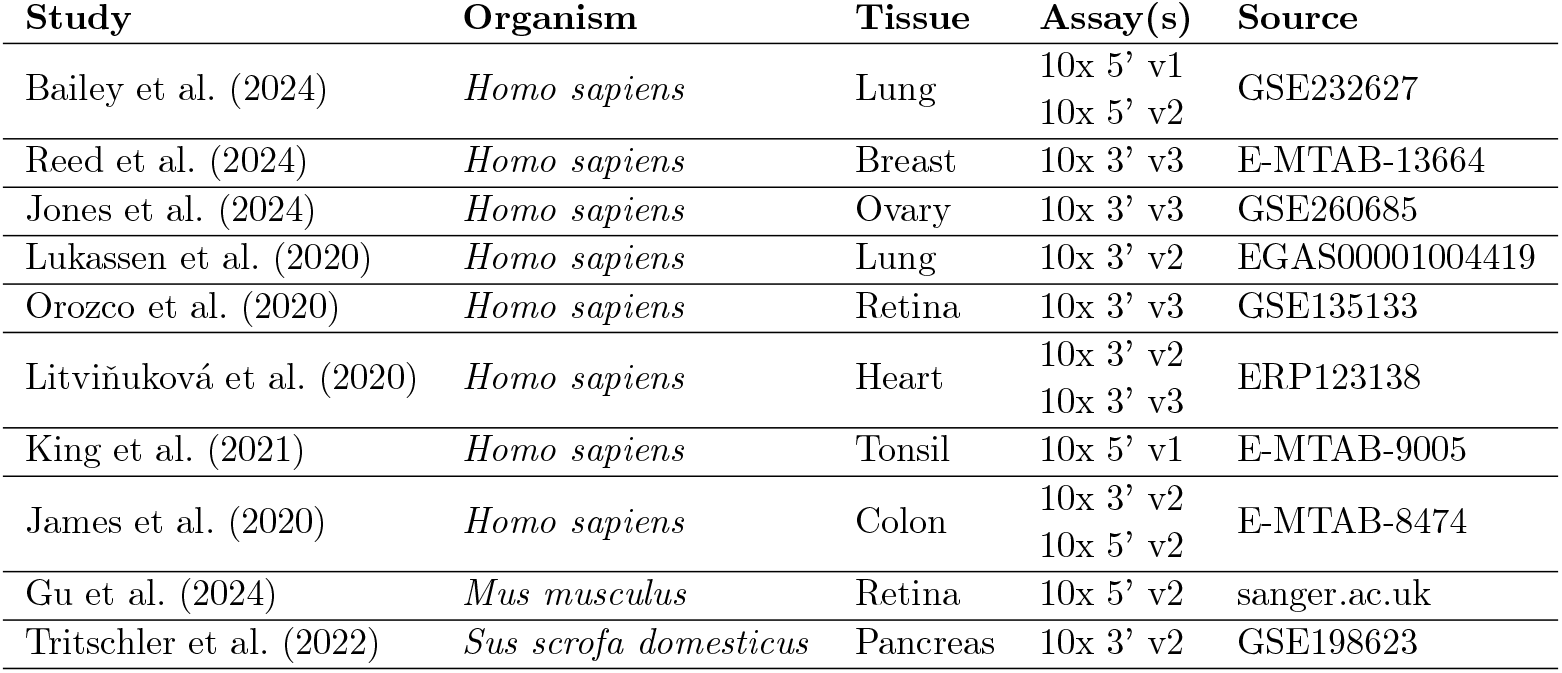
Description of scRNA-seq studies used to generate reference datasets.

The genes selected for modeling in each dataset varied between the different evaluations. For the evaluation of pairwise coexpression and low-dimensional representations, the same sets of genes were used. Genes selected for modeling were either top highly variable genes (HVGs) or genes from a predefined gene set. Some datasets had both HVGs and gene sets modeled, resulting in a total of 48 datasets being used for these evaluations. The number of genes selected in each case was between 10 and 50. HVGs were identified using the scran package (Lun et al., 2016). Gene sets were downloaded from the Kyoto Encyclopedia of Genes and Genomes (KEGG) (Kanehisa and Goto, 2000). The gene sets selected were breast cancer (hsa05224), estrogen signaling pathway (hsa04915), cardiac muscle contraction (hsa04260), and insulin secretion (hsa04911). These correspond with the tissues sequenced by Reed et al. (2024), Jones et al. (2024), Litviňuková et al. (2020), and Tritschler et al. (2022), respectively. Three of the gene sets contained more than 50 genes and were downsampled. To do so, we first constructed undirected gene-gene graphs for each gene set using graphite (Sales et al., 2012). Leiden clustering was then performed on each graph, and a cluster with 10–50 genes was chosen, yielding a “core” network. For a given dataset and gene set, genes in the gene set that were expressed by less than 2% of cells were removed.

A minimum of 500 genes is recommended for gene coexpression module identification (Morabito et al., 2023). As such, for the evaluation of gene coexpression module preservation, we needed to increase the number of genes modeled in each dataset. For each of the 38 datasets, we removed genes expressed in less than 5% of cells. We then identified the top HVGs in each dataset, and kept between 500 and 1000 genes per dataset. The distribution of cells and gene counts across reference datasets is shown in Figure S1. A description of each dataset is given in Table S1.

### 2.4 Model evaluation

Each reference scRNA-seq dataset was first split into equal sized training and testing datasets. The same set of train-test splits was used for all copula families for a given evaluation. For the evaluation of pairwise coexpression and low-dimensional representations, 50 train-test splits were used. For the evaluation of gene coexpression networks, only 10 train-test splits were used due to the longer run time of the evaluation.

Joint distribution functions were fit to each of the training datasets by first fitting marginal models (Section S1) and then copula models (Section S3). To account for sampling variability, each joint distribution was sampled 20 times to generate 20 synthetic datasets. The statistics for each of the evaluations were first averaged over the 20 synthetic datasets per train-test split, and then averaged again over all train-test splits, resulting in one statistic per copula per reference dataset. The different evaluations we performed are described below.

#### 2.4.1 Pairwise coexpression

To evaluate the ability of a copula model to capture pairwise gene coexpression patterns present in a reference dataset, we computed gene-gene association matrices using six measures of pairwise association: Pearson correlation, Spearman rank correlation, Kendall rank correlation, mutual information, biweight midcorrelation, and distance correlation. Given a synthetic dataset and a reference dataset, we computed the respective pairwise similarity matrices **M**_*s*_ and **M**_*r*_, and then computed the Frobenius norm of their difference:

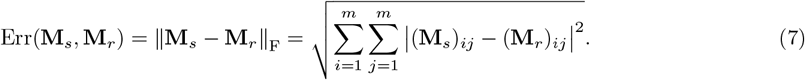

We refer to this quantity as Frobenius error.

#### 2.4.2 Low-dimensional representations

Next, we sought to evaluate how similar synthetic datasets looked to reference datasets when embedded in a low-dimensional space. Differences between the embeddings could be driven by differences in either pairwise or higher order coexpression patterns. For each reference dataset, we performed principal component analysis (PCA) and projected the dataset onto its first two principal components (PCs). We then projected each corresponding synthetic dataset onto this PC space and tested whether the differences between the embeddings were statistically distinguishable. We did so using the Fasano-Franceschini test (Fasano and Franceschini, 1987), which is a multivariate extension of the Kolmogorov-Smirnov test. The Fasano-Franceschini test has been shown to have good power for two-sample multivariate goodness-of-fit testing, especially when testing copula alternatives (Puritz et al., 2023). The null hypothesis of the two-sample Fasano-Franceschini test is that the two samples being tested were drawn from the same distribution. Since the marginal distributions for a given reference dataset are fixed, any difference in the embeddings is due to the choice of copula model. As such, we would expect larger (less significant) *p*-values for better fitting copula models.

#### 2.4.3 Gene coexpression modules

We also sought to evaluate how copula models can impact gene coexpression module identification. We reasoned that since coexpression modules are determined by analyzing gene-gene correlations, a more accurate copula model would lead to better preservation of coexpression modules in synthetic datasets. We identified gene coexpression modules using hdWGCNA (Morabito et al., 2023), which is an extension of WGCNA (Langfelder and Horvath, 2008) for scRNA-seq data.

Metacells were constructed in training and testing datasets using the MetacellsByGroups function, with the target number of metacells set to 500. We then identified gene coexpression modules in the testing datasets using the TestSoftPowers and ConstructNetwork functions. Next, we projected the modules from each testing dataset onto the corresponding simulated datasets using the ProjectModules function. We then performed module preservation testing using the ModulePreservation function with 100 permutations. We quantified module preservation using the *Z*_summary_ statistic, which is a composite statistic that combines several *Z* statistics which individually measure different aspects of module preservation. Simulations have shown this statistic to perform well at distinguishing preserved from non-preserved modules (Langfelder et al., 2011). Since the number of modules identified varied between reference datasets, we computed the mean *Z*_summary_ statistic across all modules, and used this value to compare copula models. We denote this value as 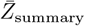.

#### 2.4.4 Run time

To evaluate the run time for fitting different copulas, we randomly downsampled the Bailey24 dataset (Bailey et al., 2024) to generate datasets with 100, 1000, 2000, 3000, 4000, or 5000 cells and 10, 20, 30, 40, or 50 genes. For each combination of the number of cells and genes, we fit Gaussian and jittered Gaussian copulas on 100 random datasets. Vine, jittered vine, ML Gaussian, and *t* copulas were fit only on three random datasets each due to their longer run times. The reported times for each copula represent the average across all datasets. Each task was run on a single CPU core of an Intel Xeon CPU (Gold 6230 @ 2.10GHz, Gold 6338 @ 2.0GHz, or Platinum 8592+ @ 1.9GHz) and was provided with a maximum of 50GB of RAM. Fitting was terminated if the wall time exceeded 10 days.

## 3 Results

### 3.1 Preservation of pairwise gene coexpression

We first evaluated the different copula models on their ability to accurately capture pairwise gene coexpression patterns present in reference datasets (see Section 2.4.1). Coexpression was quantified using six different commonly used measures of association, and copulas were evaluated on the Frobenius error between the pairwise gene association matrices of each synthetic dataset and corresponding reference dataset (Figure 2).

**Figure 2:**
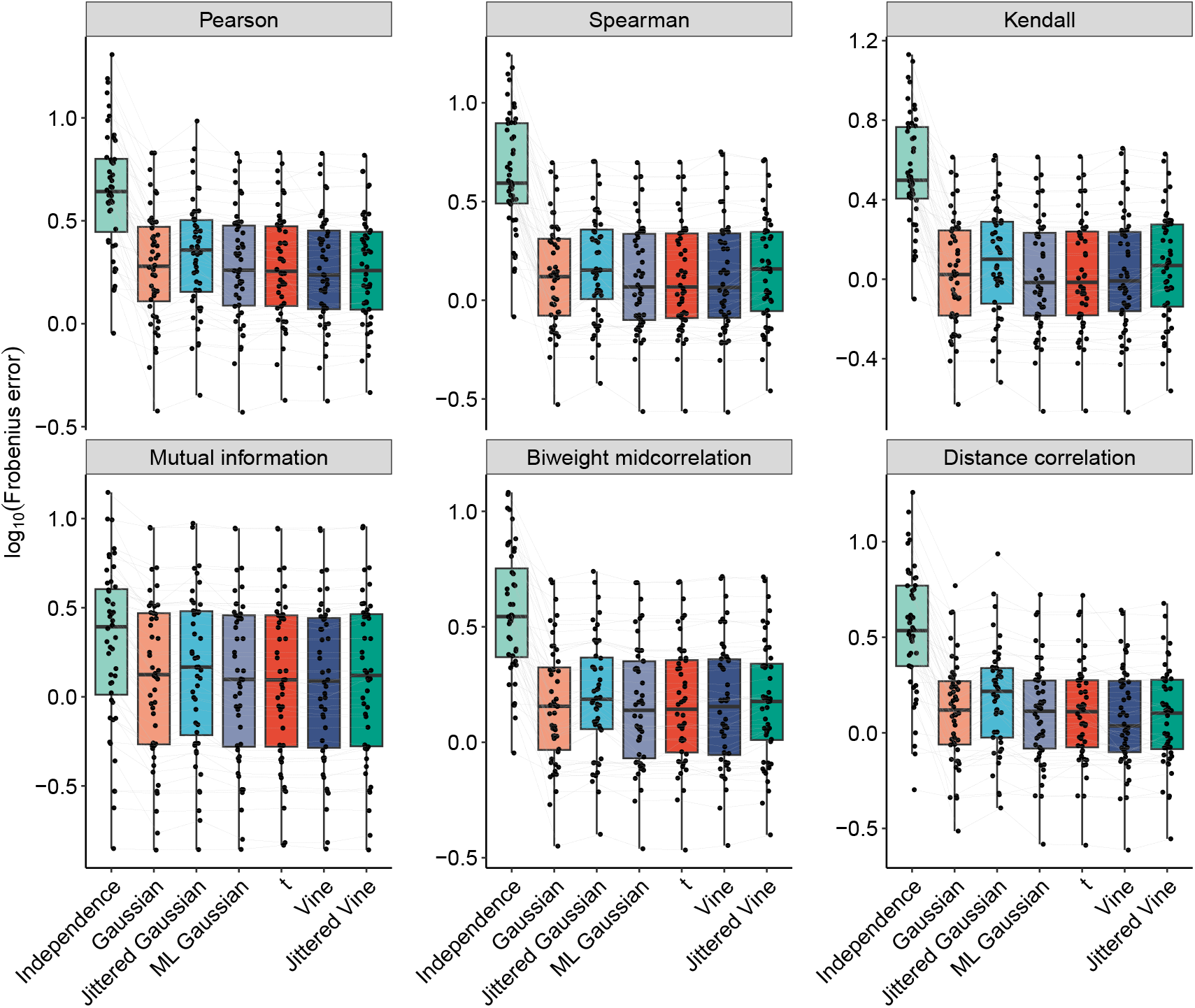
Evaluation of copula models on preserving pairwise gene associations. Comparison of copula models on their ability to preserve pairwise gene associations in reference datasets. Each point represents the average Frobenius error between gene-gene association matrices of synthetic datasets and the corresponding reference dataset. Measures of association include Pearson correlation, Spearman rank correlation, Kendall rank correlation, mutual information, biweight midcorrelation, and distance correlation. Different models of the same reference dataset are connected by a line.

For each measure of association, we tested whether the error was significantly different (paired Wilcoxon rank-sum test with FDR correction, *α <* 0.05) between each pair of copula families (excluding independence copulas). Out of the 90 pairwise comparisons, 51 were significant (see Table S2). However, many of these comparisons had extremely small effect sizes, suggesting that the differences between models may be negligible. Nevertheless, some patterns are notable: out of the 26 comparisons with an effect size larger than 0.10, 20 comparisons involve jittered Gaussian copulas, with jittered Gaussian copulas having a higher error than the other family in every comparison. Furthermore, in none of those 26 comparisons do vine, ML Gaussian, or *t* copulas perform worse than another family. We also observe that there is generally an increase in error when using jittered copulas as opposed to fitting the same type of copula (Gaussian or vine) directly to transcript counts. ML Gaussian and *t* copulas have nearly identical performance with each other and similar performance to vine copulas across all measures of association.

### 3.2 Similarity of low-dimensional embeddings

To further assess model accuracy, we evaluated how similar embeddings of synthetic datasets were to reference datasets in a low-dimensional space (see Section 2.4.2). Since the marginal distributions remain fixed for each reference dataset, any differences in embeddings are due solely to the choice of copula. These could be driven by differences in either pairwise or higher order gene coexpression patterns. For each pair of synthetic and reference datasets, we projected both onto the two-dimensional principal component space of the reference dataset. The similarity of the two embeddings was then evaluated using the Fasano-Franceschini test, which evaluates the null hypothesis that two samples were drawn from the same underlying distribution. Larger *p*-values therefore indicate that the embeddings are harder to statistically distinguish, suggesting that the copula model more accurately captures coexpression patterns in the reference dataset.

As before, jittered Gaussian copulas perform worst, tending to have the smallest (worst) *p*-values besides the independence copula (Figure 3). Vine and jittered vine copulas perform best, with vine copulas generally having slightly larger (better) *p*-values that jittered vine copulas on the same reference datasets. Gaussian, ML Gaussian, and *t* copulas all perform similarly. These results are consistent when the number of principal components used is increased Figure S2).

**Figure 3:**
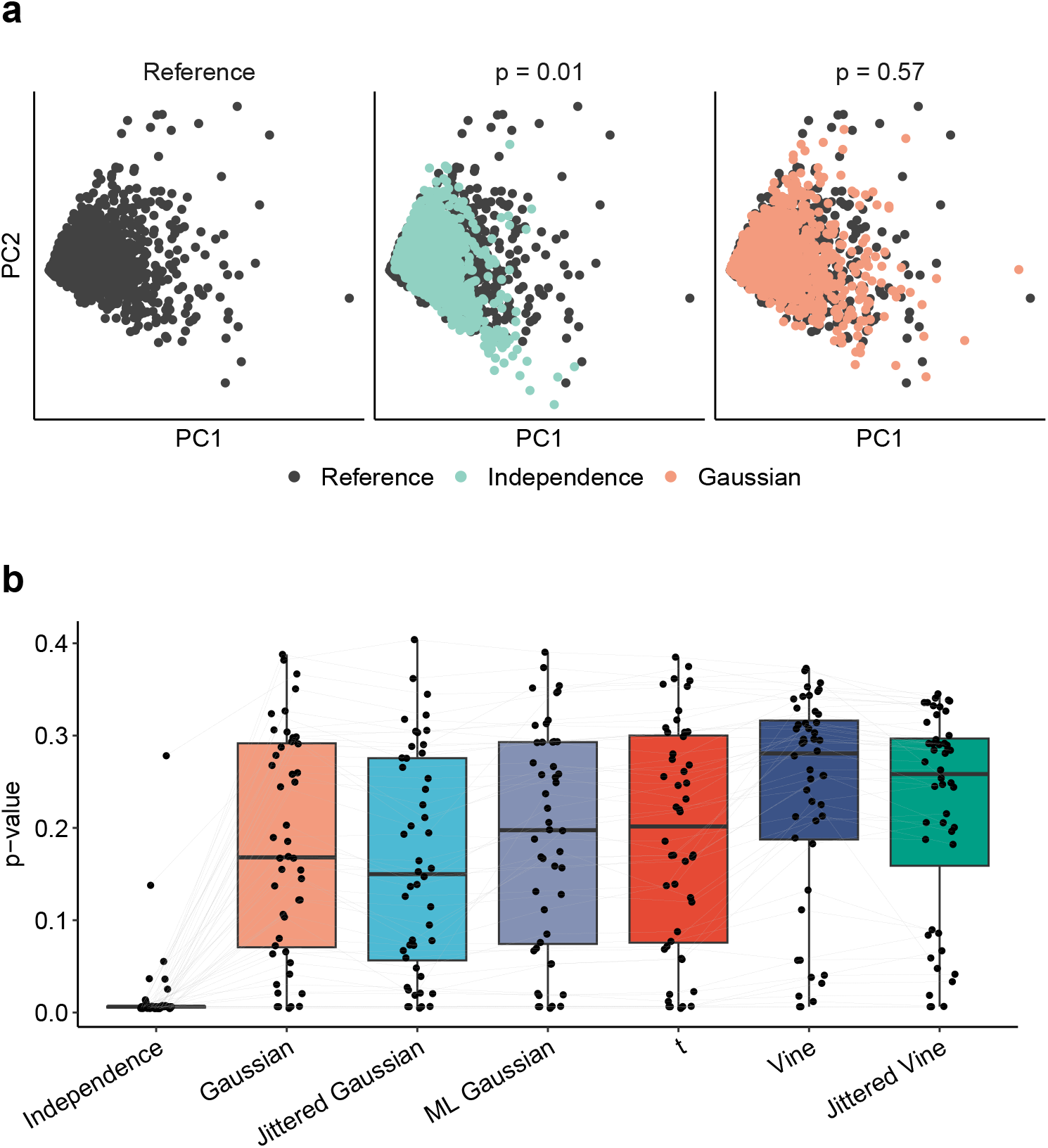
Evaluation of copula models using principal component embeddings. (a) Example of synthetic datasets generated from two different copula models of the same reference dataset embedded in the two-dimensional principal component (PC) space of the reference dataset. The *p*-value for the Fasano-Franceschini test between the embeddings is shown above each plot. (b) Evaluation of PC embeddings of synthetic datasets generated from different copula models. Each point represents the *p*-value from the Fasano-Franceschini test comparing the two-dimensional embeddings of a synthetic dataset and the corresponding reference dataset in the PC space of the reference dataset. A larger *p*-value indicates that the two embeddings are harder to statistically distinguish. Different models of the same reference dataset are connected by a line.

### 3.3 Gene coexpression network preservation

In the previous two evaluations, we examined the similarity of synthetic and reference count matrices. Next, we sought to evaluate copula models on a downstream task. A common step in the analysis of RNA-seq data is gene coexpression network analysis. In the typical methodology, networks are constructed with nodes representing genes, and the edges connecting the nodes being weighted by some measure of pairwise gene correlation. A clustering algorithm is then applied to identify groups of strongly coexpressed genes called coexpression modules (Lemoine et al., 2021). Since the choice of copula determines the coexpression patterns in synthetic datasets generated from a copula model, we reasoned that the choice of copula would impact the similarity of coexpression modules between reference and synthetic datasets.

We performed gene coexpression network analysis using hdWGCNA (Morabito et al., 2023), which is an extension of the popular tool WGCNA (Langfelder and Horvath, 2008) designed specifically for scRNA-seq data (Figure 4a). Preservation of coexpression modules was quantified using the statistic *Z*_summary_, which is a composite statistic evaluating various aspects of module preservation. The higher the value of *Z*_summary_, the better coexpression modules were preserved. Since the number of modules identified in reference datasets varied, we compared copulas by averaging *Z*_summary_ across all modules in each reference dataset (Figure 4b). We denote this value as 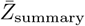.

**Figure 4:**
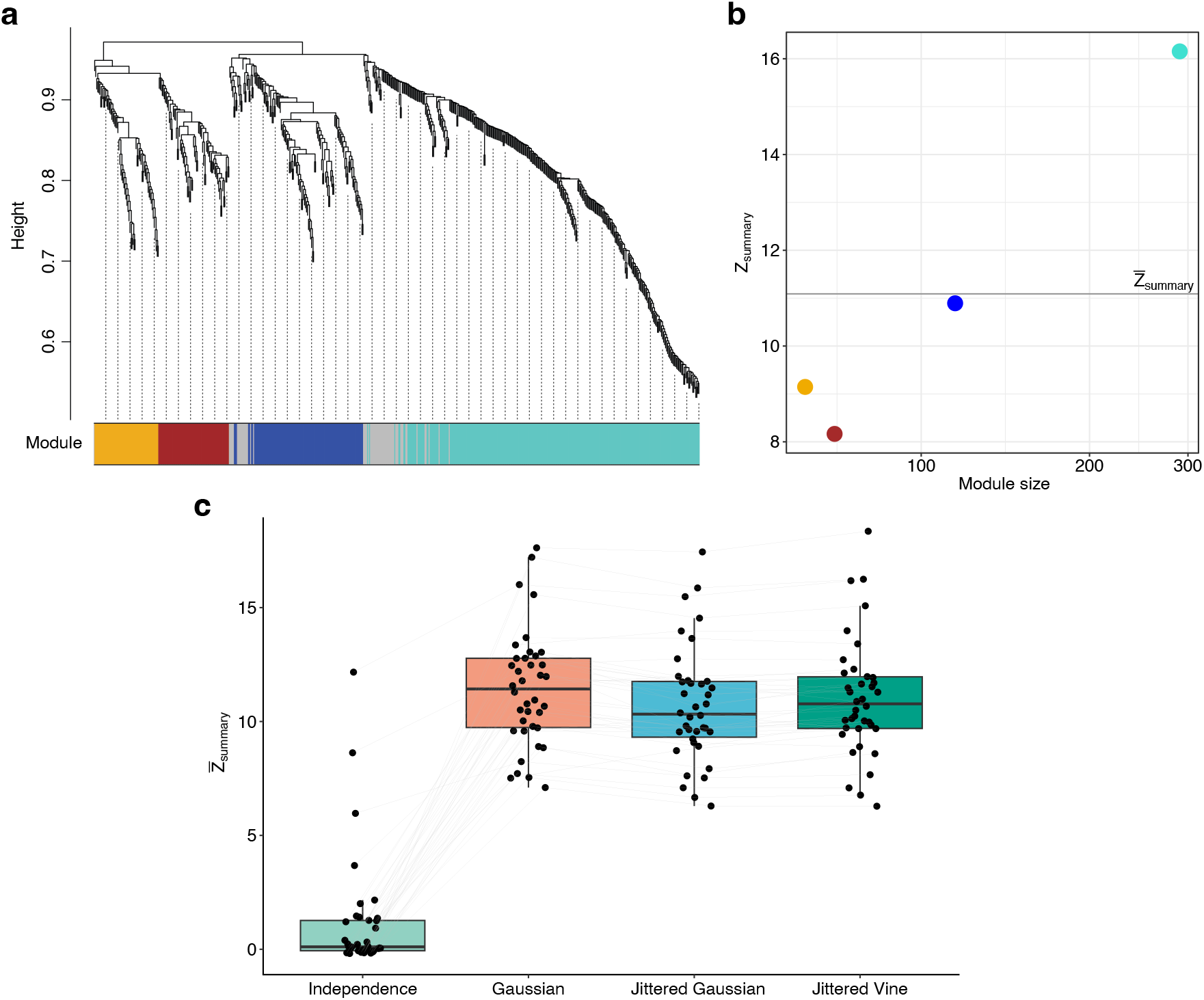
Influence of copulas on gene coexpression module preservation. (a) hdWGCNA cluster dendrogram and module assignment for genes in a reference dataset of human CD8^+^ T-cells. Module assignments are labeled by color underneath the dendrogram. Genes unassigned to a module are labeled in gray. (b) The module preservation statistic *Z*_summary_ for the four modules identified in (a), plotted as a function of module size. The *x*-axis is on a logarithmic scale. The statistic 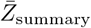 is indicated by a gray line. (c) Preservation of gene coexpression modules between reference datasets and synthetic datasets generated using different copula models. Each point represents the 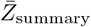 statistic for a single synthetic dataset. Different models of the same reference dataset are connected by a line.

As a minimum of 500 genes is recommended for analysis (Morabito et al., 2023), we increased the number of genes in each reference dataset to between 500 and 1000. Gaussian, jittered Gaussian, and jittered vine copulas were evaluated on how well coexpression modules in reference datasets were preserved in synthetic datasets. As before, the independence copula was included as a baseline. Due to their long run times, ML Gaussian, *t*, and vine copulas could not be fit on datasets of this size.

Gaussian copulas show the best performance (Figure 4c), having significantly higher 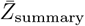 statistics (paired Wilcoxon rank-sum test with FDR correction) than both jittered Gaussian (*p* = 5.09 *×* 10^−6^) and jittered vine copulas (*p* = 2.54 *×* 10^−3^). Jittered vine copulas also outperform jittered Gaussian copulas (*p* = 2.80 *×* 10^−4^).

### 3.4 Run time

Finally, we evaluated the copulas on the time required to fit them. We found the run times to be quite disparate (Figure 5, Figure S3). Gaussian and jittered Gaussian copulas only require computing a sample correlation matrix. As such, they are the quickest to fit and have similar run times to each other. All other copula families require maximum likelihood estimation of some form, and therefore require substantially more time to fit. The times required to fit vine and jittered vine copulas scale roughly the same with respect to the number of cells and genes in the dataset, although vine copulas generally take several orders of magnitude longer to fit than jittered vine copulas on the same dataset. ML Gaussian and *t* copulas take the longest, and outside of the smallest datasets, they required more than 10 days to fit on a single core (Intel Xeon CPU @ 1.9–2.1 GHz, 50GB RAM).

**Figure 5:**
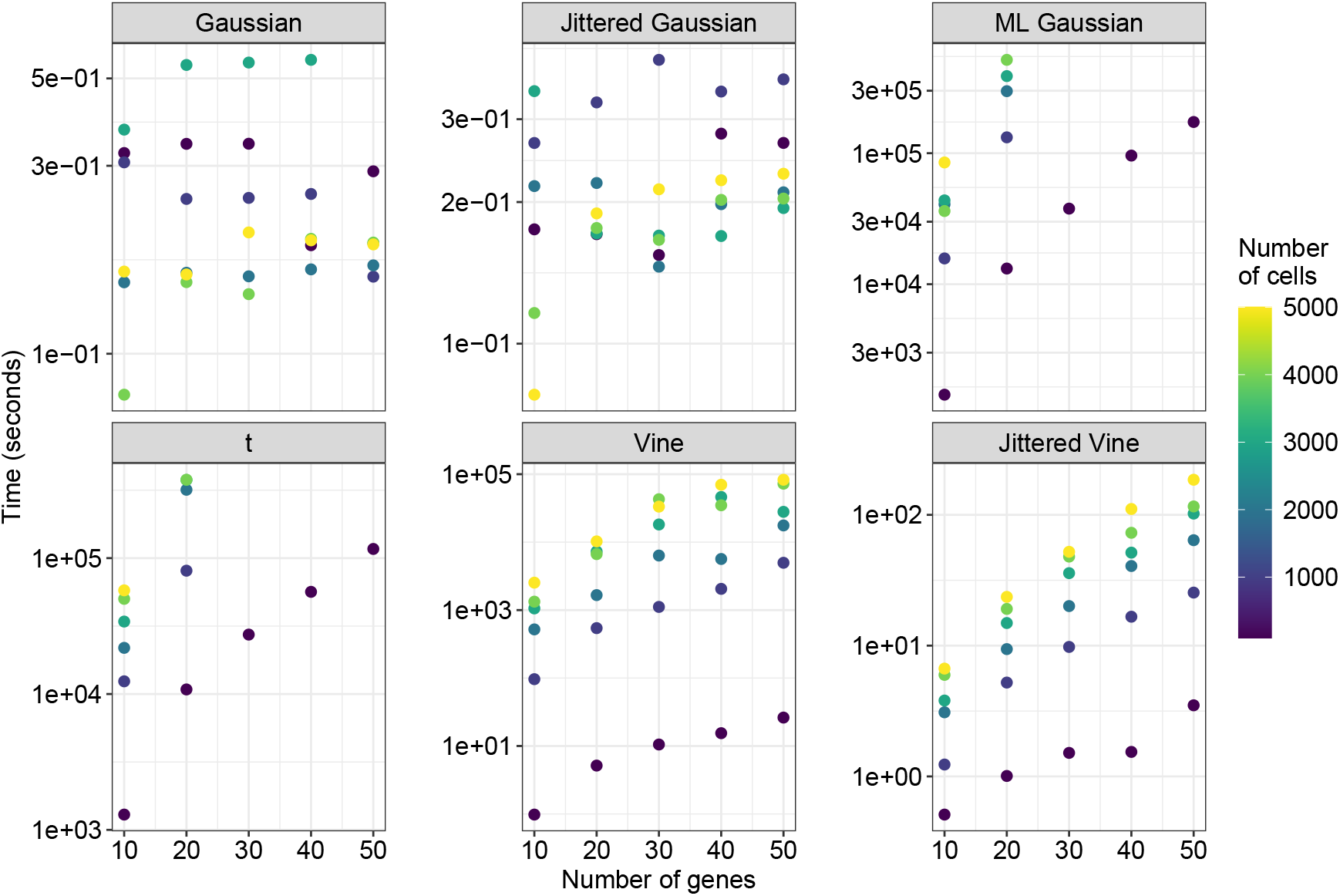
Time to fit copula models. Copula models were fit on datasets with between 100 and 5000 cells, and between 10 and 50 genes. The *y*-axes are on a logarithmic scale.

## 4 Discussion

Statistical models of scRNA-seq data form the basis of many methods for analyzing scRNA-seq data. As scRNA-seq becomes an increasingly integral tool in cell-level studies, it is has become critical to develop accurate models. Statistical models of scRNA-seq data must model not only transcript counts of individual genes, but also the patterns of coexpression between genes. Copula-based modeling has emerged as a popular approach to coexpression modeling, although no studies have evaluated the performance of different copula models in doing so. In this study, we systematically evaluated six copula models on their ability to accurately and efficiently capture coexpression patterns in scRNA-seq data.

Our results suggest that there is no benefit in using a more flexible copula model as opposed to a simple Gaussian copula with a sample correlation matrix estimator. Using ML Gaussian or *t* copulas provides only a negligible increase in accuracy at the cost of a generally impractical amount of time required to fit them. Jittered Gaussian copulas had a markedly worse performance while offering no increase in speed. The evaluations comparing the accuracy of Gaussian and jittered vine copulas were mixed with no clear top performer. However, fitting jittered vine copulas takes several orders or magnitude longer. The only family which does show a nontrivial improvement in performance are vine copulas. However, as with the other families requiring maximum likelihood estimation using raw transcript counts, the use of vine copulas is hampered by the time required to fit them. On small datasets (*<* 10k cells, *<* 100 genes), it is feasible to fit them. On modeling tasks that require high accuracy, we would recommend vine copulas if sufficient computing resources are available. However, scRNA-seq studies now commonly sequence tens of thousands if not millions of cells, and if a large number of genes need to be modeled, vine copulas will generally be infeasible to fit. Strategies to better balance accuracy and speed for vine copulas could therefore be a possible avenue for future research.

## 5 Acknowledgments

This research was supported in part through the computational resources and staff contributions provided for the Quest high performance computing facility at Northwestern University which is jointly supported by the Office of the Provost, the Office for Research, and Northwestern University Information Technology.

This research was supported in part through the computational resources and staff contributions provided by the Genomics Compute Cluster which is jointly supported by the Feinberg School of Medicine, the Center for Genetic Medicine, and Feinberg’s Department of Biochemistry and Molecular Genetics, the Office of the Provost, the Office for Research, and Northwestern Information Technology. The Genomics Compute Cluster is part of Quest, Northwestern University’s high performance computing facility, with the purpose to advance research in genomics.

## 6 Funding

This work was supported by NSF grant DMS-1764421, Simons Foundation grant 597491, and NIH grant R01AG068579.

## 7 Code availability

All code used to generate the results and figures in this manuscript is available at https://github.com/cpuritz/scrna_copula_modeling.

## 8 Data availability

All datasets used in this study are publicly available. All scRNA-seq datasets except for Bailey24 are available in AnnData format from CZ CELLxGENE (https://cellxgene.cziscience.com/datasets). The Bailey24 dataset (Bailey et al., 2024) is available under GEO accession GSE232627. The Reed24 dataset (Reed et al., 2024) is available under EMBL accession E-MTAB-13664. The Jones24 (Jones et al., 2024) dataset is available under GEO accession GSE260685. The Lukassen20 dataset (Lukassen et al., 2020) is available under EGA accession EGAS00001004419. The Orozco20 dataset (Orozco et al., 2020) is available under GEO accession GSE135133. The Litvinukova20 dataset (Litviňuková et al., 2020) is available under ENA accession ERP123138. The King21 dataset (King et al., 2021) is available under EMBL accession EMTAB-9005. The James20 dataset (James et al., 2020) is available under EMBL accession E-MTAB-8474. The Gu24 dataset (Gu et al., 2024) is available from https://treg-gut-niches.cellgeni.sanger.ac.uk. The Tritschler22 dataset (Tritschler et al., 2022) is available under GEO accession GSE198623. All gene sets are available from KEGG.

## Supplementary Material

### S1 Marginal distributions

We model marginal distributions empirically to avoid degrading copula parameter estimates in the case of marginal misspecification. Specifically, for a sample 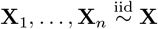, we model the *i*th marginal distribution *F*_*i*_ using the estimator

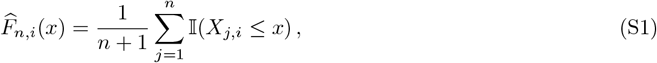

where 𝕀 is the indicator function. Note that this estimator is *n/*(*n*+1) times the typical empirical distribution function. This asymptotically negligible rescaling is used to avoid issues with evaluating quantile functions on the boundaries of the unit hypercube (Genest et al., 1995).

### S2 Jittered copulas

Let **X** be a *d*-dimensional random variable with joint distribution function *H* and marginal distribution functions 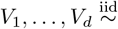 Unif(0, 1) be random variables that are independent of **X**. Define the *d*-dimensional random variable **U** by

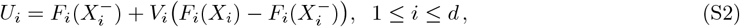

where 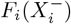 denotes the left-limit of *F*_*i*_ at *X*_*i*_. The margins of **U** are uniformly distributed on the unit interval, and the joint distribution function of **U** is a copula itself (Nešlehová, 2007). This copula, which we denote as 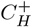, is a copula of *H* (Nešlehová, 2007). *If H* were continuous, then it would be the unique copula of *H*. In literature, 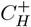 is often referred to as the multilinear extension copula, as it can be constructed by linear interpolation on the images of the marginal distribution functions (Genest et al., 2013).

We refer to the process of constructing **U** from **X** as jittering. Other names for this technique in literature include distributional transformation (e.g. Song et al., 2024) and continuous extension (e.g. Nešlehová, 2007). Jittering before performing inference has two supposed benefits. First, it enables the use of copula parameter inference methods for continuous distributions, which tend to be much faster than those for discontinuous distributions (Panagiotelis et al., 2012). Second, it resolves the identifiability issue, since the copula of **U** is unique.

However, jittering also introduce new complications. The process of jittering introduces additional variability through the realizations of the uniform noise terms *V*_*i*_. In this work, we only use a single jitter. To account for this added variability, likelihood equations could be modified to be averaged over a large number of jitters (Nikoloulopoulos, 2013). However, this will lead to longer computational times, nullifying some of benefit gained by jittering in the first place. Furthermore, the estimators obtained from jittering may not be unbiased. Nikoloulopoulos (2013) found that the maximum likelihood estimators for Gaussian copula correlation matrices obtained using jittering were biased, and often underestimated the true strength of dependence. Increasing the number of jitters reduced the variability of the estimators but did not decrease the bias.

### S3 Copula inference

Let **X** be a count-valued random variable with marginal distribution functions **F** = (*F*_1_, …, *F*_*d*_). Let **X**_1_, …, **X**_*n*_ be independent and identically distributed samples of **X**. We performed copula inference semiparametrically using empirical estimators of the marginal distribution functions. Under mild regularity conditions, the maximum likelihood estimators of the copula parameters are consistent when using empirical estimators of count-valued marginal distribution functions (Nasri and Remillard, 2023). Here, we describe the inference procedure for each class of copulas.

#### S3.1 Gaussian & jittered Gaussian copulas

For the Gaussian copula, the estimator of the correlation matrix is the sample correlation matrix of the normal-transformed pseudo-observations

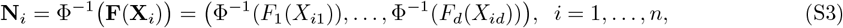

where **Φ**^−1^ is the standard normal quantile function. For the jittered Gaussian copula, jittered pseudo-observations (Equation S2) are used instead of pseudo-observations.

#### S3.2 ML Gaussian copula

Suppose that the copula of **X** is a Gaussian copula with correlation matrix **R**. The mass function of **X** is given by

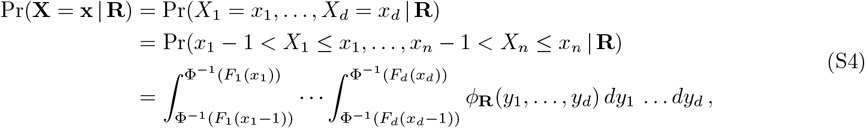

where **Φ**^−1^ is the standard normal quantile function and *ϕ*_**R**_ is the density function of a multivariate Gaussian random variable with mean **0** and covariance matrix **R** (Panagiotelis et al., 2012). The average log-likelihood of the samples **X**_1_, …, **X**_*n*_ is given by

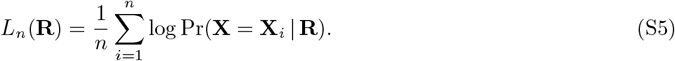

As there is no closed-form expression for the mass function, we compute it numerically using the method of Botev (2016) implemented in the TruncatedNormal package (Botev and Belzile, 2024). We used 1000 Monte Carlo simulations as we found this to provide a good balance between time and accuracy.

Fitting a Gaussian copula to the samples **X**_1_, …, **X**_*n*_ requires estimating the correlation matrix **R**. A reasonable initial estimator is 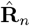, the sample correlation matrix of **N**_1_, …, **N**_*n*_, where **N**_*i*_ = **Φ**^−1^(**F**(**X**_*i*_)). Note however that 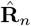 is not the maximum likelihood estimator (Hernández et al., 2014). To the best of our knowledge, no closed form solution exists for the maximum likelihood estimator, and thus we need to perform numerical maximization. To do so, we need to reparametrize the likelihood function since updating elements of **R** directly during optimization may produce a matrix which is no longer a valid correlation matrix (i.e., a symmetric positive semi-definite matrix with ones on the diagonal). We can parametrize **R** as a length *d*(*d* − 1)*/*2 vector using the approach of Bhat and Mondal (2021), which we describe below. Note that we will assume that **R** is positive definite. If **R** was a singular positive semi-definite matrix, the resulting distribution would be valid but degenerate.

The unique Cholesky factor of **R** can be written as

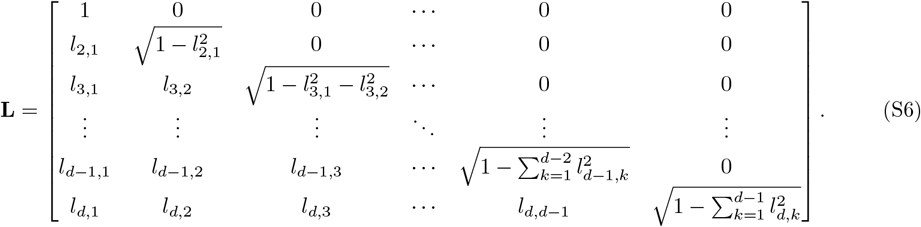

Suppose we parametrize the lower triangular elements of **L** as

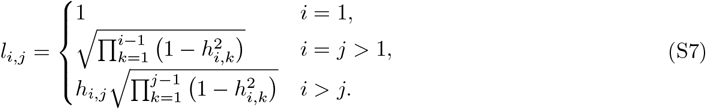

This is a valid parametrization if we take **H** = (*h*_*i,j*_) to be the *d × d* lower triangular matrix with *h*_*i,i*_ = 1 and *h*_*i*,1_ = *l*_*i*,1_ for all 1 ≤ *i* ≤ *d*, and

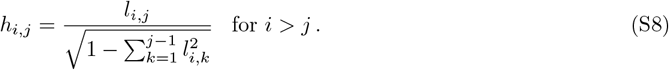

This parametrization of **L** is useful since it ensures the diagonal elements of **L** will remain positive and real during the search process. Since the diagonal elements of **H** are all 1, we can uniquely encode **H** as a vector **v**_**H**_ ∈ (−1, 1)^*d*(*d*−1)*/*2^ whose elements are the lower triangular elements of **H** in column-major order. The elements of **v**_**H**_ can then be mapped to the real line using the bijection *f*: (−1, 1) → ℝ defined by

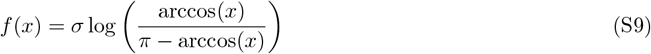

with inverse

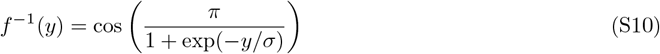

where *σ >* 0 is a scale parameter. This parametrization constrains the search procedure to the space of symmetric positive-definite matrices. Numerical experiments performed by Bhat and Mondal (2021) suggest that when this parametrization of the likelihood function is maximized using the BFGS algorithm, a scale parameter of *σ* = 1.2904 provides a good balance between convergence time and accuracy, and thus this is the value we use. Maximization of Equation S5 was performed numerically using L-BFGS-B optimization (Byrd et al., 1995).

#### S3.3 *t* copula

Suppose that the copula of **X** is a *t* copula with degrees of freedom *ν >* 0 and correlation matrix **R**. The mass function of **X** is given by

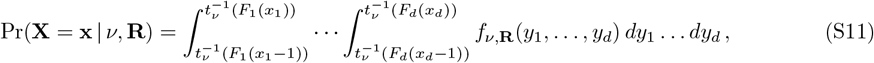

where 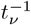 is the univariate *t* quantile function, and *f*_*ν*,**R**_ is the density function of a multivariate *t* random variable with mean **0**, covariance matrix **R**, and degrees of freedom *ν* (Panagiotelis et al., 2012). The average log-likelihood of the samples **X**_1_, …, **X**_*n*_ is then given by

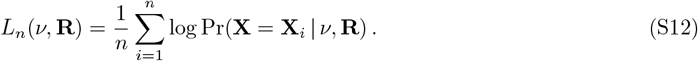

As with the Gaussian copula, there is no closed-form expression for the mass function, and we compute it numerically using the method of Botev and L’Ecuyer (2015) implemented in the TruncatedNormal package (Botev and Belzile, 2024) with 1000 Monte Carlo simulations.

While maximization of Equation S12 could be performed jointly over *ν* and **R**, we instead took a two-stage approach to reduce compute time by taking advantage of the correlation matrix maximum likelihood estimates already computed for Gaussian copulas. The two-stage maximization procedure (Zeevi and Mashal, 2002) is as follows:

1. The *t* copula correlation matrix **R** is estimated by the maximum likelihood estimator 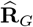 of the correlation matrix using a Gaussian copula model.
2. The degrees of freedom *ν* is estimated as

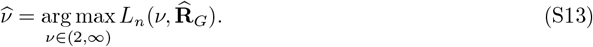

By restricting *ν* to be larger than 2, we avoid obtaining a distribution with an undefined covariance matrix. We solved Equation S13 numerically using L-BFGS-B optimization (Byrd et al., 1995).

#### S3.4 Vine & jittered vine copulas

Vine copulas were fit using the vinecop function in the rvinecopulib package (Nagler and Vatter, 2023). The pair copula family set included the independence copula, the Gaussian copula, and the four available one-parameter bivariate Archimedean copulas (Clayton, Gumbel, Frank, and Joe). Parameter estimation was performed using maximum likelihood estimation (par_method = “mle”). The Akaike information criterion (AIC) was used for family selection (selcrit = “aic”). For vine copulas fit directly to transcript counts, var types was set to “d”. For jittered vine copulas, jittered pseudo-observations (Equation S2) were passed to vinecop instead of pseudo-observations, and var types was set to “c”. The default settings of vinecop were used otherwise.

### S4 Computational details

All results were generated using R 4.4.0 (R Core Team, 2024) on Northwestern’s Quest high-performance computing cluster, which consists of nodes equipped with Intel Emerald Rapids Xeon Platinum 8592+ @ 1.9GHz, Intel Ice Lake Xeon Gold 6338 @ 2.0GHz, or Intel Cascade Lake Xeon Gold 6230 @ 2.10GHz processors. Gene sets were processed using igraph 2.1.4 (Csárdi and Nepusz, 2006), graphite 1.52.0 (Sales et al., 2012), org.Hs.eg.db 3.20.0 (Carlson, 2024), and AnnotationDbi 1.68.0 (Pag`es et al., 2024). Multivariate elliptical distribution functions were evaluated numerically using TruncatedNormal 2.3 (Botev and Belzile, 2024). Vine copulas were fit and sampled from using rvinecopulib 0.7.2.1.0 (Nagler and Vatter, 2023). All other copulas were sampled from using copula 1.1-6 (Hofert et al., 2024). Likelihood maximization was performed using the optim function in the stats package (R Core Team, 2024) or the optimParallel function from optimParallel 1.0-2 (Gerber and Furrer, 2019). Midweight bicorrelation was computed using the bicor function from WGCNA 1.73 (Langfelder and Horvath, 2008). Distance correlation was computed using the dcor function from energy 1.7-12 (Rizzo and Szekely, 2024). Fasano-Franceschini tests were performed using fasano.franceschini.test 2.2.2 (Puritz et al., 2023). Gene coexpression module analysis was performed using hdWGCNA 0.4.07 (Morabito et al., 2023). Cohen’s *d* was calculated using effsize 0.8.1 (Torchiano, 2020). Wilcoxon rank-sum tests were performed using coin 1.4-3 (Hothorn et al., 2006).

Figures were generated using R 4.4.3 (R Core Team, 2024) on macOS using dplyr 1.1.4 (Wickham et al., 2023), tidyr 1.3.1 (Wickham et al., 2024), ggplot2 3.5.2 (Wickham, 2016), ggsci 3.2.0 (Xiao, 2024), ggsignif 0.6.4 (Constantin and Patil, 2021), scales 1.4.0 (Wickham et al., 2025), reshape2 1.4.4 (Wickham, 2007), and patchwork 1.3.2 (Pedersen, 2024).

**Table S1:** Description of scRNA-seq datasets used as references for simulation.

| Study | Dataset | Genes | Cells | Description |
| --- | --- | --- | --- | --- |
| Bailey24 | bailey24-rpra04-moam | 17 | 1030 | Profibrotic monocyte-derived alveolar macrophages from donor RPRA04. Subset to top HVGs. |
|  | bailey24-rpra09-tram | 12/500 | 2668 | Tissue-resident alveolar macrophages from donor RPRA09. Subset to top HVGs/top HVGs. |
|  | bailey24-rpra24-tram | 15/600 | 1672 | Tissue-resident alveolar macrophages from donor RPRA24. Subset to top HVGs/top HVGs. |
|  | bailey24-rpra29-dc2 | 35 | 602 | Type II conventional dendritic cells from donor RPRA29. Subset to top HVGs. |
|  | bailey24-rpra20-cd8t | 40/500 | 2285 | CD8<sup>+</sup> T cells from donor RPRA20. Subset to top HVGs/top HVGs. |
|  | bailey24-rpra30-cd4t | 32/825 | 3040 | CD4<sup>+</sup> T cells from donor RPRA30. Subset to top HVGs/top HVGs. |
| Reed24 | reed24-donor6-myoepithelial | 30/20/775 | 1510 | Basal myoepithelial cells from donor 6. Subset to top HVGs/hsa05224 gene set/top HVGs. |
|  | reed24-donor18-myoepithelial | 36/21/675 | 4389 | Basal myoepithelial cells from donor 18. Subset to top HVGs/hsa05224 gene set/top HVGs. |
|  | reed24-donor19-myoepithelial | 45/20/725 | 2340 | Basal myoepithelial cells from donor 19. Subset to top HVGs/hsa05224 gene set/top HVGs. |
|  | reed24-donor26-myoepithelial | 50/19 | 1056 | Basal myoepithelial cells from donor 26. Subset to top HVGs/hsa05224 gene set. |
| Jones24 | jones24-donor3-stromal | 25/22 | 750 | Ovarian stromal cells from donor 3. Subset to top HVGs/hsa04915 gene set. |
|  | jones24-donor4-stromal | 20/23/750 | 1500 | Ovarian stromal cells from donor 4. Subset to top HVGs/hsa04915 gene set/top HVGs. |
|  | jones24-donor5-stromal | 15/24/900 | 2400 | Ovarian stromal cells from donor 5. Subset to top HVGs/hsa04915 gene set/top HVGs. |
| Lukassen20 | lukassen20-9JQK55ng-monocyte | 32 | 1163 | Monocytes from donor 9JQK55ng. Subset to top HVGs. |
|  | lukassen20-A9LCTZng-monocyte | 10 | 1062 | Monocytes from donor A9LCTZng. Subset to top HVGs. |
|  | lukassen20-QZY9VQng-monocyte | 14 | 955 | Monocytes from donor QZY9VQng. Subset to top HVGs. |
| Orozco20 | orozco20-109373-mueller | 20/625 | 2017 | Müller cells from donor 109373. Subset to top HVGs/top HVGs. |
|  | orozco20-109829-mueller | 12 | 1140 | Müller cells from donor 109829. Subset to top HVGs. |
|  | orozco20-110814-mueller | 42 | 530 | Müller cells from donor 110814. Subset to top HVGs. |
| Orozco20 | orozco20-120974-mueller | 10 | 1427 | Müller cells from donor 120974. Subset to top HVGs. |
| Litvinukova20 | litvinukova20-d3-myocyte | 21/1000 | 2656 | Atrial cardiac myocytes from donor D3. Cells were sequenced using 10x 3' v2. Subset to hsa04260 gene set/top HVGs. |
|  | litvinukova20-d6-myocyte | 21/525 | 2116 | Atrial cardiac myocytes from donor D6. Cells were sequenced using 10x 3' v2. Subset to hsa04260 gene set/top HVGs. |
|  | litvinukova20-h6-myocyte | 21 | 1400 | Atrial cardiac myocytes from donor H6. Cells were sequenced using 10x 3' v3. Subset to hsa04260 gene set. |
|  | litvinukova20-h7-myocyte | 21/575 | 3469 | Atrial cardiac myocytes from donor H7. Cells were sequenced using 10x 3' v3. Subset to hsa04260 gene set/top HVGs. |
| King21 | king21-BCP5-naiveB | 16/950 | 2262 | Naive B cells from donor BCP5. Subset to top HVGs/top HVGs. |
|  | king21-BCP6-naiveB | 20 | 1106 | Naive B cells from donor BCP6. Subset to top HVGs. |
|  | king21-BCP8-naiveB | 24 | 477 | Naive B cells from donor BCP8. Subset to top HVGs. |
| James20 | james20-290b-memB | 10 | 1392 | Memory B cells from donor 290b. Cells in this subset were sequenced using 10x 3' v2. Subset to top HVGs. |
|  | james20-302c-memB | 36/800 | 1696 | Memory B cells from donor 302c. Cells in this subset were sequenced using 10x 3' v2. Subset to top HVGs/top HVGs. |
|  | james20-390c-memB | 48 | 450 | Memory B cells from donor 390c. Cells in this subset were sequenced using 10x 5' v2. Subset to top HVGs. |
|  | james20-290b-iga | 15/700 | 5024 | IgA<sup>+</sup> plasma cells from donor 290b. Cells in this subset were sequenced using 10x 3' v2. Subset to top HVGs/top HVGs. |
|  | james20-298c-iga | 26/850 | 4736 | IgA<sup>+</sup> plasma cells from donor 298c. Cells in this subset were sequenced using 10x 3' v2. Subset to top HVGstop HVGs. |
| Gu24 | gu24-pooled-mp | 10 | 589 | Macrophages from all donors pooled. Subset to top HVGs. |
|  | gu24-pooled-dc | 46 | 912 | Conventional dendritic cells from all donors pooled. Subset to top HVGs. |
|  | gu24-pooled-plasma | 28 | 1423 | Plasma cells from all donors pooled. Subset to top HVGs. |
| Tritschler22 | tritschler22-pig1-panB | 10/27/650 | 7200 | Pancreatic β-cells from pig 1. Genes were mapped to human orthologs. Subset to top HVGs/hsa04911 gene set/top HVGs. |
| Tritschler22 | tritschler22-pig2-panA | 30/25 | 877 | Pancreatic α-cells from pig 2. Genes were mapped to human orthologs. Subset to top HVGs/hsa04911 gene set. |
| | tritschler22-pig2-panB | 20/27/875 | 6000 | Pancreatic β-cells from pig 2. Genes were mapped to human orthologs. Subset to top HVGs/hsa04911 gene set/top HVGs. |

**Table S2:** Output of paired Wilcoxon rank-sum tests on the Frobenius errors of pairwise gene association matrices. Independence copulas were excluded from testing. The columns family1 and family2 indicate the copula families compared. measure indicates the measure of association used (dcor, distance correlation; bicor, biweight midcorrelation; MI, mutual information). pval records the test *p*-value. stat records the test statistic. padj records the FDR-adjusted *p*-value. eff records the absolute value of the effect size (Cohen’s *d* for paired samples).

| family1 | family2 | measure | pval | stat | padj | eff |
| --- | --- | --- | --- | --- | --- | --- |
| Jittered Gaussian | Vine | dcor | 7.49423e-08 | 4.91289 | 2.69792e-07 | 0.32265 |
| Jittered Gaussian | Jittered Vine | dcor | 1.68754e-11 | 5.71290 | 1.51879e-10 | 0.30620 |
| Jittered Gaussian | Vine | pearson | 1.11130e-05 | 4.16416 | 2.70316e-05 | 0.24387 |
| Jittered Gaussian | Jittered Vine | pearson | 9.32816e-06 | 4.19493 | 2.39867e-05 | 0.24081 |
| Jittered Gaussian | ML Gaussian | kendall | 9.02943e-10 | 5.39494 | 4.06325e-09 | 0.21006 |
| Jittered Gaussian | ML Gaussian | dcor | 8.08399e-10 | 5.40520 | 3.82926e-09 | 0.20630 |
| Jittered Gaussian | t | dcor | 2.24517e-07 | 4.76929 | 7.21661e-07 | 0.20401 |
| Jittered Gaussian | t | kendall | 1.75889e-08 | 5.08725 | 7.19547e-08 | 0.20078 |
| Jittered Gaussian | ML Gaussian | spearman | 3.19146e-09 | 5.27186 | 1.36777e-08 | 0.19285 |
| Jittered Gaussian | t | spearman | 3.85520e-08 | 4.99494 | 1.44570e-07 | 0.18567 |
| Jittered Gaussian | Gaussian | kendall | 9.94760e-14 | 5.96931 | 4.05009e-12 | 0.18539 |
| Jittered Gaussian | ML Gaussian | pearson | 3.22331e-10 | 5.48725 | 1.81311e-09 | 0.17121 |
| Jittered Gaussian | t | pearson | 1.65420e-07 | 4.81032 | 5.51399e-07 | 0.16899 |
| Jittered Gaussian | Gaussian | spearman | 3.90799e-13 | 5.91803 | 8.79297e-12 | 0.16673 |
| Jittered Gaussian | Gaussian | dcor | 1.35003e-13 | 5.95905 | 4.05009e-12 | 0.14937 |
| Jittered Gaussian | Gaussian | pearson | 7.10543e-15 | 6.03085 | 6.39488e-13 | 0.14723 |
| Jittered Gaussian | Vine | kendall | 3.04041e-05 | 3.97954 | 7.01633e-05 | 0.13656 |
| Gaussian | Vine | dcor | 2.61503e-02 | 2.21541 | 3.67739e-02 | 0.13202 |
| Jittered Gaussian | Vine | spearman | 5.43088e-05 | 3.86672 | 1.16376e-04 | 0.12262 |
| Jittered Gaussian | ML Gaussian | bicor | 3.77700e-07 | 4.69750 | 1.13310e-06 | 0.11700 |
| Jittered Vine | ML Gaussian | kendall | 1.92311e-08 | 5.07699 | 7.52520e-08 | 0.11500 |
| Gaussian | Jittered Vine | dcor | 2.22522e-01 | 1.23079 | 2.86099e-01 | 0.11197 |
| Jittered Vine | ML Gaussian | spearman | 1.31010e-07 | 4.84109 | 4.53497e-07 | 0.11045 |
| Jittered Vine | t | kendall | 7.70744e-07 | 4.59493 | 2.10203e-06 | 0.10636 |
| Jittered Gaussian | Gaussian | bicor | 2.28297e-11 | 5.69238 | 1.86789e-10 | 0.10610 |
| Jittered Vine | t | spearman | 1.73832e-06 | 4.47185 | 4.60143e-06 | 0.10386 |
| Jittered Gaussian | Vine | MI | 5.38876e-11 | 5.63084 | 3.46420e-10 | 0.09896 |
| Jittered Gaussian | t | bicor | 1.39676e-04 | 3.67184 | 2.79351e-04 | 0.09855 |
| t | Vine | dcor | 2.54415e-02 | 2.22567 | 3.63450e-02 | 0.09352 |
| Jittered Gaussian | Jittered Vine | kendall | 3.26133e-07 | 4.71801 | 1.01214e-06 | 0.09213 |
| Jittered Gaussian | t | MI | 1.23563e-11 | 5.73341 | 1.23563e-10 | 0.08474 |
| Jittered Gaussian | Vine | bicor | 1.44321e-03 | 3.11799 | 2.59778e-03 | 0.08438 |
| Jittered Gaussian | ML Gaussian | MI | 3.17613e-12 | 5.81546 | 5.71276e-11 | 0.08252 |
| Jittered Vine | Gaussian | kendall | 5.41434e-12 | 5.78469 | 6.96129e-11 | 0.08194 |
| ML Gaussian | Vine | dcor | 4.98621e-01 | 0.68719 | 5.54023e-01 | 0.08167 |
| Jittered Vine | Gaussian | spearman | 9.23208e-11 | 5.58982 | 5.53925e-10 | 0.07730 |
| Jittered Gaussian | Jittered Vine | spearman | 1.04856e-05 | 4.17441 | 2.62139e-05 | 0.07682 |
| t | Jittered Vine | dcor | 7.25823e-01 | 0.35898 | 7.67584e-01 | 0.07245 |
| Gaussian | Vine | pearson | 6.07569e-01 | 0.52308 | 6.58809e-01 | 0.06969 |
| Jittered Vine | Gaussian | pearson | 7.33469e-01 | 0.34872 | 7.67584e-01 | 0.06502 |
| Jittered Gaussian | Jittered Vine | MI | 1.23563e-11 | 5.73341 | 1.23563e-10 | 0.06299 |
| Jittered Vine | ML Gaussian | dcor | 8.34973e-01 | 0.21539 | 8.44355e-01 | 0.06074 |
| t | Vine | pearson | 4.12411e-01 | 0.83078 | 4.82039e-01 | 0.06052 |
| Jittered Vine | ML Gaussian | bicor | 3.97025e-05 | 3.92826 | 8.93305e-05 | 0.05837 |
| Jittered Gaussian | Gaussian | MI | 3.80851e-12 | 5.80520 | 5.71276e-11 | 0.05754 |
| Jittered Gaussian | Jittered Vine | bicor | 3.64891e-03 | 2.86158 | 5.97094e-03 | 0.05655 |
| Jittered Vine | t | pearson | 9.55433e-01 | 0.06154 | 9.55433e-01 | 0.05578 |
| Vine | ML Gaussian | pearson | 8.27045e-01 | 0.22564 | 8.44355e-01 | 0.05388 |

Table S2: Output of paired Wilcoxon rank-sum tests on the Frobenius errors of pairwise gene association matrices.
| **family1** | **family2** | **measure** | **pval** | **stat** | **padj** | **eff** |
| --- | --- | --- | --- | --- | --- | --- |
| Gaussian | ML Gaussian | dcor | 4.76965e-04 | 3.39492 | 8.94310e-04 | 0.04940 |
| Jittered Vine | ML Gaussian | pearson | 4.24167e-01 | 0.81027 | 4.89424e-01 | 0.04907 |
| Vine | ML Gaussian | kendall | 8.57044e-05 | 3.77441 | 1.75304e-04 | 0.04875 |
| Jittered Vine | Vine | spearman | 1.96106e-03 | 3.03594 | 3.46070e-03 | 0.04808 |
| Vine | ML Gaussian | spearman | 6.70167e-04 | 3.31286 | 1.23092e-03 | 0.04604 |
| Jittered Vine | Vine | kendall | 2.27797e-03 | 2.99491 | 3.86825e-03 | 0.04590 |
| Jittered Vine | Gaussian | bicor | 6.24475e-07 | 4.62570 | 1.75634e-06 | 0.04551 |
| Vine | t | kendall | 1.56634e-02 | 2.40003 | 2.34951e-02 | 0.04183 |
| Gaussian | Vine | MI | 6.46175e-10 | 5.42571 | 3.42092e-09 | 0.04143 |
| Jittered Vine | t | bicor | 4.05445e-03 | 2.83081 | 6.51609e-03 | 0.04077 |
| Vine | t | spearman | 2.47490e-02 | 2.23593 | 3.59260e-02 | 0.04028 |
| Gaussian | t | dcor | 6.74349e-03 | 2.67696 | 1.04640e-02 | 0.04015 |
| Jittered Vine | Vine | MI | 7.23091e-10 | 5.41546 | 3.61545e-09 | 0.03452 |
| Vine | Gaussian | kendall | 5.65996e-02 | 1.90772 | 7.83687e-02 | 0.03150 |
| Jittered Vine | Vine | bicor | 3.16446e-03 | 2.90260 | 5.27411e-03 | 0.02853 |
| Gaussian | t | MI | 4.07709e-11 | 5.65136 | 3.05782e-10 | 0.02793 |
| Vine | Gaussian | spearman | 1.44971e-01 | 1.46669 | 1.94737e-01 | 0.02746 |
| Gaussian | ML Gaussian | MI | 4.69100e-11 | 5.64110 | 3.24762e-10 | 0.02564 |
| Vine | ML Gaussian | bicor | 1.12344e-02 | 2.51285 | 1.71373e-02 | 0.02483 |
| Gaussian | ML Gaussian | spearman | 3.51217e-01 | 0.94360 | 4.27156e-01 | 0.02458 |
| Jittered Vine | Vine | dcor | 1.62900e-01 | 1.40515 | 2.15603e-01 | 0.02329 |
| Gaussian | ML Gaussian | kendall | 3.40731e-01 | 0.96412 | 4.20079e-01 | 0.02277 |
| Jittered Vine | t | MI | 5.04453e-07 | 4.65647 | 1.46454e-06 | 0.02172 |
| Jittered Vine | ML Gaussian | MI | 2.13359e-04 | 3.57953 | 4.17442e-04 | 0.01965 |
| t | ML Gaussian | bicor | 6.01878e-05 | 3.84620 | 1.25974e-04 | 0.01886 |
| Vine | Gaussian | bicor | 2.77433e-01 | 1.09745 | 3.46791e-01 | 0.01792 |
| ML Gaussian | Vine | MI | 1.48076e-05 | 4.11287 | 3.50705e-05 | 0.01737 |
| Gaussian | t | spearman | 4.66798e-01 | 0.73847 | 5.25148e-01 | 0.01665 |
| Gaussian | ML Gaussian | pearson | 1.69227e-01 | 1.38463 | 2.20731e-01 | 0.01599 |
| t | Vine | MI | 4.37371e-04 | 3.41543 | 8.37518e-04 | 0.01504 |
| Gaussian | t | kendall | 5.79375e-01 | 0.56411 | 6.35899e-01 | 0.01240 |
| t | ML Gaussian | dcor | 3.72783e-01 | 0.90258 | 4.41454e-01 | 0.01043 |
| t | ML Gaussian | kendall | 5.15726e-05 | 3.87697 | 1.13208e-04 | 0.01027 |
| Gaussian | t | pearson | 6.95515e-01 | 0.40001 | 7.45194e-01 | 0.01011 |
| Gaussian | ML Gaussian | bicor | 2.77433e-01 | 1.09745 | 3.46791e-01 | 0.00971 |
| Vine | t | bicor | 4.48248e-01 | 0.76924 | 5.10663e-01 | 0.00936 |
| Gaussian | t | bicor | 7.95513e-01 | 0.26667 | 8.22944e-01 | 0.00916 |
| t | ML Gaussian | spearman | 2.03634e-03 | 3.02568 | 3.52443e-03 | 0.00793 |
| t | ML Gaussian | pearson | 3.67318e-01 | 0.91283 | 4.40781e-01 | 0.00600 |
| Jittered Vine | Vine | pearson | 7.33057e-02 | 1.79490 | 9.99623e-02 | 0.00512 |
| Gaussian | Jittered Vine | MI | 2.40726e-02 | 2.24618 | 3.55170e-02 | 0.00446 |
| ML Gaussian | t | MI | 6.74349e-03 | 2.67696 | 1.04640e-02 | 0.00239 |

**Figure S1:**
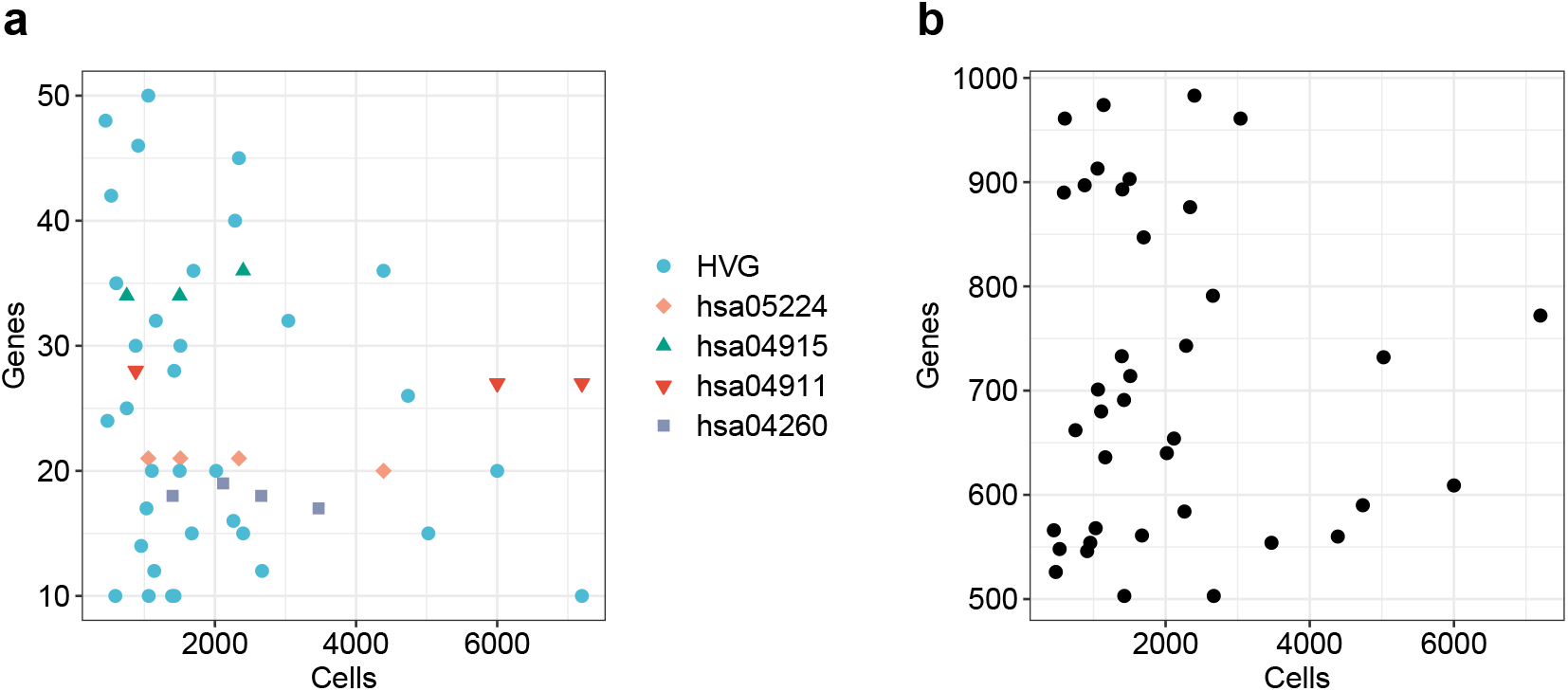
(a) Distribution of the number of cells and genes in reference scRNA-seq datasets used for pairwise coexpression and low-dimensional embedding evaluations. Genes for each dataset were either highly variable genes (HVG) or selected from the specified KEGG gene sets. (b) Distribution of the number of cells and genes in reference scRNA-seq datasets used for gene coexpression module evaluation.

**Figure S2:**
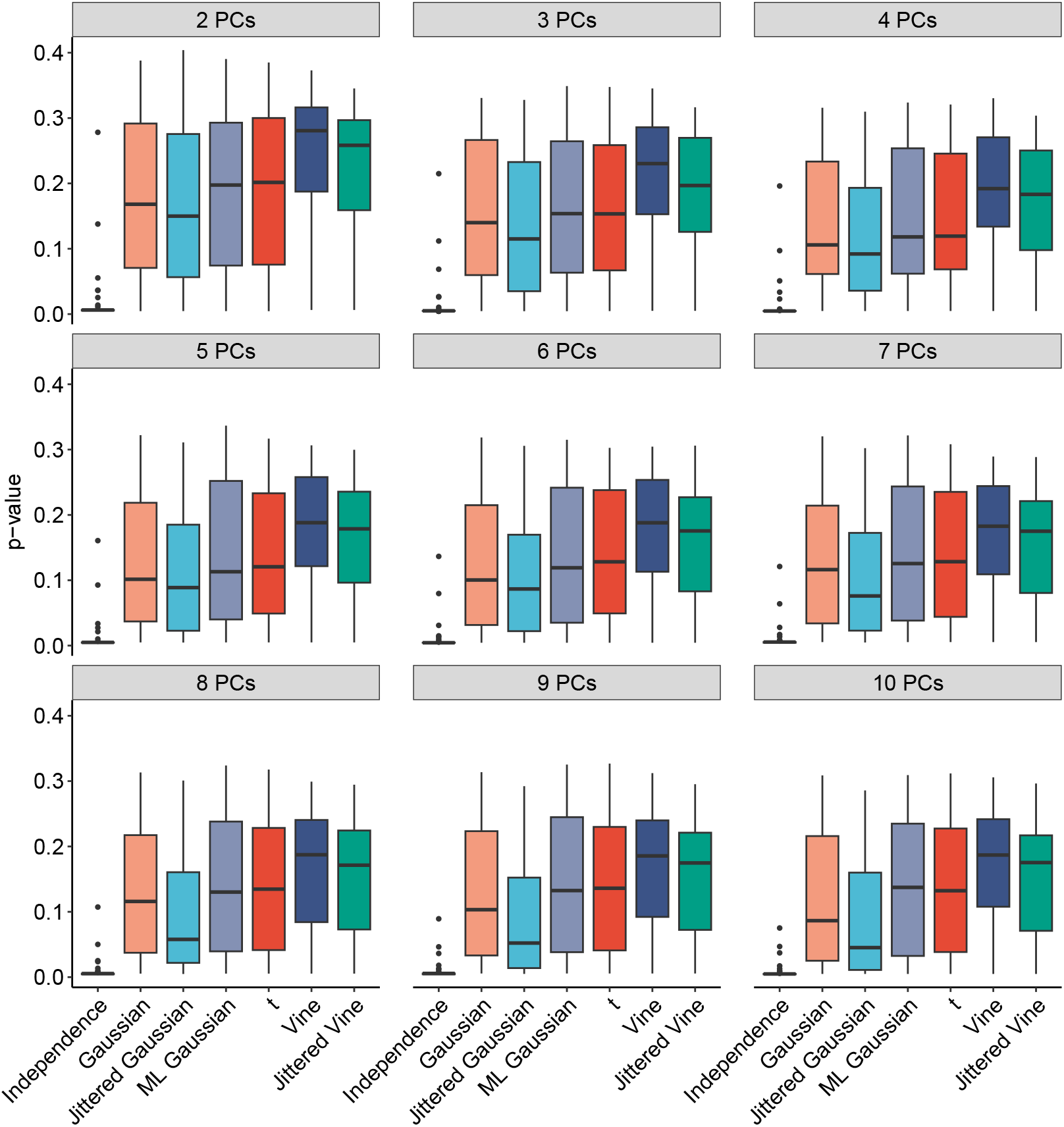
Evaluation of copula models using principal component (PC) embeddings with varying numbers of PCs. Each point represents the *p*-value from the Fasano-Franceschini test comparing the embeddings of a synthetic dataset and the corresponding reference dataset in the PC space of the reference dataset. A larger *p*-value indicates that the two embeddings are harder to statistically distinguish. The number of PCs retained for the embeddings is shown above each panel.

**Figure S3:**
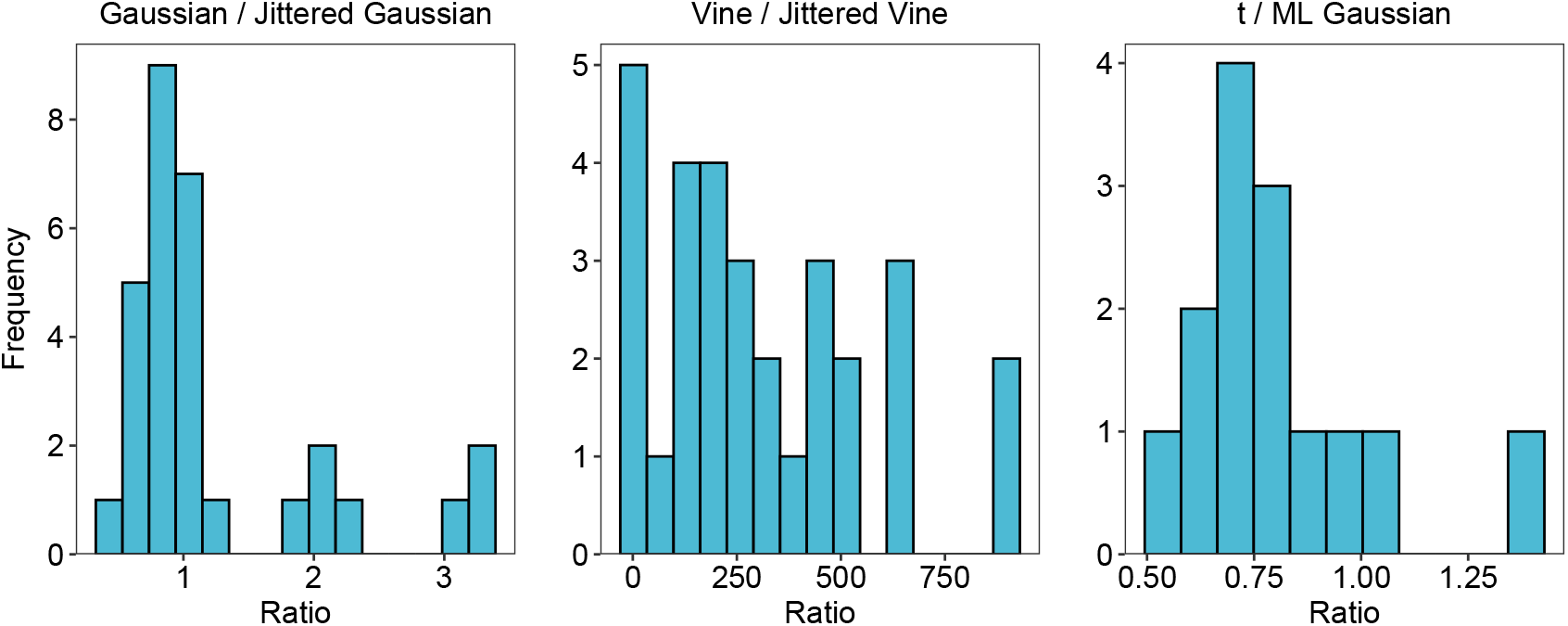
Ratio of times to fit different copula models on datasets of the same size. The times here are the same as reported in Figure 5. The ratio represented in each histogram is shown in the respective plot title.

## Footnotes

1 Discrete covariates can be modeled by fitting copulas separately to each distinct value of the covariate.

## Notes

### Competing Interest Statement

The authors have declared no competing interest.

## References

A. T. Assefa, J. Vandesompele, and O. Thas. SPsimSeq: semi-parametric simulation of bulk and single-cell RNA-sequencing data. Bioinformatics, 36(10):3276–3278, 2020. doi: 10.1093/bioinformatics/btaa105.

J. I. Bailey, C. H. Puritz, K. J. Senkow, N. S. Markov, E. Diaz, E. Jonasson, Z. Yu, S. Swaminathan, Z. Lu, S. Fenske, R. A. Grant, H. Abdala-Valencia, R. J. Mylvaganam, A. Ludwig, J. Miller, R. I. Cumming, R. M. Tighe, K. M. Gowdy, R. Kalhan, M. Jain, A. Bharat, C. Kurihara, R. San Jose Estepar, R. San Jose Estepar, G. R. Washko, A. Shilatifard, J. I. Sznajder, K. M. Ridge, G. R. S. Budinger, R. Braun, A. V. Misharin, and M. A. Sala. Profibrotic monocyte-derived alveolar macrophages are expanded in patients with persistent respiratory symptoms and radiographic abnormalities after COVID-19. Nature Immunology, 25(11):2097–2109, 2024. doi: 10.1038/s41590-024-01975-x.

C. R. Bhat and A. Mondal. On the almost exact-equivalence of the radial and spherical unconstrained Cholesky-based parameterization methods for correlation matrices. Technical report, Department of Civil, Architectural and Environmental Engineering, The University of Texas at Austin, 2021. URL https://www.caee.utexas.edu/prof/bhat/ABSTRACTS/Cholesky_parameterization.pdf.

Z. Botev and L. Belzile. TruncatedNormal: Truncated Multivariate Normal and Student Distributions, 2024. URL https://CRAN.R-project.org/package=TruncatedNormal. R package version 2.3.

Z. I. Botev. The normal law under linear restrictions: simulation and estimation via minimax tilting. Journal of the Royal Statistical Society Series B: Statistical Methodology, 79(1):125–148, 2016. doi: 10.1111/rssb.12162.

Z. I. Botev and P. L’Ecuyer. Efficient probability estimation and simulation of the truncated multivariate Student-t distribution. In 2015 Winter Simulation Conference (WSC), pages 380–391, 2015. doi: 10.1109/WSC.2015.7408180.

R. Braun, L. Cope, and G. Parmigiani. Identifying differential correlation in gene/pathway combinations. BMC bioinformatics, 9(1):488, 2008.

R. H. Byrd, P. Lu, J. Nocedal, and C. Zhu. A limited memory algorithm for bound constrained optimization. SIAM Journal on Scientific Computing, 16(5):1190–1208, 1995. doi: 10.1137/0916069.

M. Carlson. org.Hs.eg.db: Genome wide annotation for Human, 2024. URL https://bioconductor.org/packages/org.Hs.eg.db. R package version 3.20.0.

A.-E. Constantin and I. Patil. ggsignif: R package for displaying significance brackets for ‘ggplot2’. PsyArxiv, 2021. doi: 10.31234/osf.io/7awm6.

G. Csárdi and T. Nepusz. The igraph software package for complex network research. InterJournal, Complex Systems:1695, 2006. URL https://igraph.org.

C. Czado and T. Nagler. Vine copula based modeling. Annual Review of Statistics and Its Application, 9 (1):453–477, 2022. doi: 10.1146/annurev-statistics-040220-101153.

G. Fasano and A. Franceschini. A multidimensional version of the Kolmogorov-Smirnov test. Monthly Notices of the Royal Astronomical Society, 225(1):155–170, 1987. doi: 10.1093/mnras/225.1.155.

C. Fuetterer, G. Schollmeyer, and T. Augustin. Constructing simulation data with dependency structure for unreliable single-cell RNA-sequencing data using copulas. In International Symposium on Imprecise Probabilities: Theories and Applications, pages 216–224. Proceedings of Machine Learning Research, 2019. URL https://proceedings.mlr.press/v103/fuetterer19a/fuetterer19a.pdf.

C. Gaiteri, Y. Ding, B. French, G. C. Tseng, and E. Sibille. Beyond modules and hubs: the potential of gene coexpression networks for investigating molecular mechanisms of complex brain disorders. Genes, Brain and Behavior, 13(1):13–24, 2014. doi: 10.1111/gbb.12106.

C. Genest, K. Ghoudi, and L.-P. Rivest. A semiparametric estimation procedure of dependence parameters in multivariate families of distributions. Biometrika, 82(3):543–552, 1995. doi: 10.1093/biomet/82.3.543.

C. Genest, J.G. Nešlehová, and B. Rémillard. On the estimation of Spearman’s rho and related tests of independence for possibly discontinuous multivariate data. Journal of Multivariate Analysis, 117:214–228, 2013. doi: 10.1016/j.jmva.2013.02.007.

C. Genest, O. Okhrin, and T. Bodnar. Copula modeling from Abe Sklar to the present day. Journal of Multivariate Analysis, 201:105278, 2024. doi: 10.1016/j.jmva.2023.105278.

F. Gerber and R. Furrer. optimparallel: An R package providing a parallel version of the L-BFGS-B optimization method. The R Journal, 11(1):352–358, 2019. doi: 10.32614/RJ-2019-030.

Y. Gu, R. Bartolomé-Casado, C. Xu, A. Bertocchi, A. Janney, C. Heuberger, C. F. Pearson, S. A. Teichmann, E. E. Thornton, and F. Powrie. Immune microniches shape intestinal Treg function. Nature, 628(8009):854–862, 2024. doi: 10.1038/s41586-024-07251-0.

L. Hernández, J. Tejero, and J. Vinuesa. Maximum likelihood estimation of the correlation parameters for elliptical copulas. arXiv, 2014. doi: 10.48550/arXiv.1412.6316.

Y.-Y. Ho, L. Cope, M. Dettling, and G. Parmigiani. Statistical methods for identifying differentially expressed gene combinations. In Gene Function Analysis, pages 171–191. Springer, 2007.

M. Hofert, I. Kojadinovic, M. Maechler, and J. Yan. copula: Multivariate Dependence with Copulas, 2024. URL https://CRAN.R-project.org/package=copula. R package version 1.1-4.

T. Hothorn, K. Hornik, M. A. van de Wiel, and A. Zeileis. A Lego system for conditional inference. The American Statistician, 60(3):257–263, 2006. doi: 10.1198/000313006X118430.

K. R. James, T. Gomes, R. Elmentaite, N. Kumar, E. L. Gulliver, H. W. King, M. D. Stares, B. R. Bareham, J. R. Ferdinand, V. N. Petrova, K. Polański, S. C. Forster, L. B. Jarvis, O. Suchanek, S. Howlett, L. K. James, J. L. Jones, K. B. Meyer, M. R. Clatworthy, K. Saeb-Parsy, T. D. Lawley, and S. A. Teichmann. Distinct microbial and immune niches of the human colon. Nature Immunology, 21(3):343–353, 2020. doi: 10.1038/s41590-020-0602-z.

A. S. K. Jones, D. F. Hannum, J. H. Machlin, A. Tan, Q. Ma, N. D. Ulrich, Y. Shen, M. Ciarelli, V. Padmanabhan, E. E. Marsh, S. Hammoud, J. Z. Li, and A. Shikanov. Cellular atlas of the human ovary using morphologically guided spatial transcriptomics and single-cell sequencing. Science Advances, 10 (14):eadm7506, 2024. doi: 10.1126/sciadv.adm7506.

M. Kanehisa and S. Goto. KEGG: Kyoto Encyclopedia of Genes and Genomes. Nucleic Acids Research, 28 (1):27–30, 2000. doi: 10.1093/nar/28.1.27.

H. W. King, N. Orban, J. C. Riches, A. J. Clear, G. Warnes, S. A. Teichmann, and L. K. James. Single-cell analysis of human B cell maturation predicts how antibody class switching shapes selection dynamics. Science Immunology, 6(56):eabe6291, 2021. doi: 10.1126/sciimmunol.abe6291.

P. Langfelder and S. Horvath. WGCNA: an R package for weighted correlation network analysis. BMC Bioinformatics, 9(1):559, 2008. doi: 10.1186/1471-2105-9-559.

P. Langfelder, R. Luo, M. C. Oldham, and S. Horvath. Is my network module preserved and reproducible? PLoS computational biology, 7(1):e1001057, 2011. doi: 10.1371/journal.pcbi.1001057.

G. G. Lemoine, M.-P. Scott-Boyer, B. Ambroise, O. Périn, and A. Droit. GWENA: gene co-expression networks analysis and extended modules characterization in a single Bioconductor package. BMC Bioinformatics, 22(1):267, 2021. doi: 10.1186/s12859-021-04179-4.

W. V. Li and J. J. Li. A statistical simulator scDesign for rational scRNA-seq experimental design. Bioinformatics, 35(14):i41–i50, 2019. doi: 10.1093/bioinformatics/btz321.

M. Litviňuková, C. Talavera-López, H. Maatz, D. Reichart, C. L. Worth, E. L. Lindberg, M. Kanda, K. Polanski, M. Heinig, M. Lee, E. R. Nadelmann, K. Roberts, L. Tuck, E. S. Fasouli, D. M. DeLaughter, B. McDonough, H. Wakimoto, J. M. Gorham, S. Samari, K. T. Mahbubani, K. Saeb-Parsy, G. Patone, J. J. Boyle, H. Zhang, H. Zhang, A. Viveiros, G. Y. Oudit, O. A. Bayraktar, J. G. Seidman, C. E. Seidman, M. Noseda, N. Hubner, and S. A. Teichmann. Cells of the adult human heart. Nature, 588(7838):466–472, 2020. doi: 10.1038/s41586-020-2797-4.

S. Lukassen, R. L. Chua, T. Trefzer, N. C. Kahn, M. A. Schneider, T. Muley, H. Winter, M. Meister, C. Veith, A. W. Boots, B. P. Hennig, M. Kreuter, C. Conrad, and R. Eils. SARS-CoV-2 receptor ACE2 and TMPRSS2 are primarily expressed in bronchial transient secretory cells. The EMBO Journal, 39(10): e105114, 2020. doi: 10.15252/embj.20105114.

A. T. L. Lun, D. J. McCarthy, and J. C. Marioni. A step-by-step workflow for low-level analysis of single-cell RNA-seq data with Bioconductor. F1000Research, 5:2122, 2016. doi: 10.12688/f1000research.9501.2.

S. Morabito, F. Reese, N. Rahimzadeh, E. Miyoshi, and V. Swarup. hdWGCNA identifies co-expression networks in high-dimensional transcriptomics data. Cell Reports Methods, 3(6), 2023. doi: 10.1016/j.crmeth.2023.100498.

T. Nagler and T. Vatter. rvinecopulib: High Performance Algorithms for Vine Copula Modeling, 2023. URL https://CRAN.R-project.org/package=rvinecopulib. R package version 0.6.3.1.1.

B. R. Nasri and B. N. Remillard. Identifiability and inference for copula-based semiparametric models for random vectors with arbitrary marginal distributions. arXiv, 2023. doi: 10.48550/arXiv.2301.13408.

R. B. Nelsen. An introduction to copulas. Springer Series in Statistics. Springer, New York, 2nd edition, 2006. ISBN 978-0-387-28659-4.

J. Nešlehová. On rank correlation measures for non-continuous random variables. Journal of Multivariate Analysis, 98(3):544–567, 2007. doi: 10.1016/j.jmva.2005.11.007.

A. K. Nikoloulopoulos. On the estimation of normal copula discrete regression models using the continuous extension and simulated likelihood. Journal of Statistical Planning and Inference, 143(11):1923–1937, 2013. doi: 10.1016/j.jspi.2013.06.015.

L. D. Orozco, H.-H. Chen, C. Cox, K. J. Katschke, R. Arceo, C. Espiritu, P. Caplazi, S. S. Nghiem, Y.-J. Chen, Z. Modrusan, A. Dressen, L. D. Goldstein, C. Clarke, T. Bhangale, B. Yaspan, M. Jeanne, M. J. Townsend, M. van Lookeren Campagne, and J. A. Hackney. Integration of eQTL and a single-cell atlas in the human eye identifies causal genes for age-related macular degeneration. Cell Reports, 30(4): 1246–1259.e6, 2020. doi: 10.1016/j.celrep.2019.12.082.

H. Pagés, M. Carlson, S. Falcon, and N. Li. AnnotationDbi: Manipulation of SQLite-based annotations in Bioconductor, 2024. URL https://bioconductor.org/packages/AnnotationDbi. R package version 1.68.0.

Panagiotelis, C. Czado, and H. Joe. Pair copula constructions for multivariate discrete data. Journal of the American Statistical Association, 107(499):1063–1072, 2012.

Panwar, B. J. Schmiedel, S. Liang, B. White, E. Rodriguez, K. Kalunian, A. J. McKnight, R. Soloff, G. Seumois, P. Vijayanand, and F. Ay. Multi–cell type gene coexpression network analysis reveals coordinated interferon response and cross–cell type correlations in systemic lupus erythematosus. Genome Research, 31(4):659–676, 2021. doi: 10.1101/gr.265249.120.

T. L. Pedersen. patchwork: The Composer of Plots, 2024. URL https://CRAN.R-project.org/package=patchwork. R package version 1.3.2.

C. Puritz, E. Ness-Cohn, and R. Braun. fasano.franceschini.test: An implementation of a multivariate KS test in R. The R Journal, 15(3):159–171, 2023. doi: 10.32614/RJ-2023-067.

R Core Team. R: A Language and Environment for Statistical Computing. R Foundation for Statistical Computing, Vienna, Austria, 2024. URL https://www.R-project.org/.

S. Ray, S. Lall, and S. Bandyopadhyay. CODC: a copula-based model to identify differential coexpression. npj Systems Biology and Applications, 6(1):20, 2020. doi: 10.1038/s41540-020-0137-9.

A. D. Reed, S. Pensa, A. Steif, J. Stenning, D. J. Kunz, L. J. Porter, K. Hua, P. He, A.-J. Twigger, A. J. Q. Siu, K. Kania, R. Barrow-McGee, I. Goulding, J. J. Gomm, V. Speirs, J. L. Jones, J. C. Marioni, and W. T. Khaled. A single-cell atlas enables mapping of homeostatic cellular shifts in the adult human breast. Nature Genetics, 56(4):652–662, 2024. doi: 10.1038/s41588-024-01688-9.

D. Risso, F. Perraudeau, S. Gribkova, S. Dudoit, and J.-P. Vert. A general and flexible method for signal extraction from single-cell RNA-seq data. Nature Communications, 9(1):284, 2018. doi: 10.1038/s41467-017-02554-5.

M. Rizzo and G. Szekely. energy: E-Statistics: Multivariate Inference via the Energy of Data, 2024. URL https://CRAN.R-project.org/package=energy. R package version 1.7-12.

C. Ruprecht, N. Vaid, S. Proost, S. Persson, and M. Mutwil. Beyond genomics: Studying evolution with gene coexpression networks. Trends in Plant Science, 22(4):298–307, 2017.

G. Sales, E. Calura, D. Cavalieri, and C. Romualdi. graphite - a Bioconductor package to convert pathway topology to gene network. BMC Bioinformatics, 2012. doi: 10.1186/1471-2105-13-20.

H. Sarkar, U. Chitra, J. Gold, and B. J. Raphael. A count-based model for delineating cell–cell interactions in spatial transcriptomics data. Bioinformatics, 40(Supplement 1):i481–i489, 2024. doi: 10.1093/bioinformatics/btae219.

A. Sklar. Fonctions de répartition án dimensions et leurs marges. Publ. Inst. Statist. Univ. Paris, 8:229–231, 1959.

D. Song, Q. Wang, G. Yan, T. Liu, T. Sun, and J. J. Li. scDesign3 generates realistic in silico data for multimodal single-cell and spatial omics. Nature Biotechnology, 42(2):247–252, 2024. doi: 10.1038/s41587-023-01772-1.

J. M. Stuart, E. Segal, D. Koller, and S. K. Kim. A gene-coexpression network for global discovery of conserved genetic modules. Science, 302(5643):249–255, 2003. doi: 10.1126/science.1087447.

T. Sun, D. Song, W. V. Li, and J. J. Li. scDesign2: a transparent simulator that generates high-fidelity single-cell gene expression count data with gene correlations captured. Genome Biology, 22(1):163, 2021. doi: 10.1186/s13059-021-02367-2.

J. Tian, J. Wang, and K. Roeder. ESCO: single cell expression simulation incorporating gene co-expression. Bioinformatics, 37(16):2374–2381, 2021. doi: 10.1093/bioinformatics/btab116.

M. Torchiano. effsize: Efficient Effect Size Computation, 2020. URL https://CRAN.R-project.org/package=effsize. R package version 0.8.1.

S. Tritschler, M. Thomas, A. Böttcher, B. Ludwig, J. Schmid, U. Schubert, E. Kemter, E. Wolf, H. Lickert, and F. J. Theis. A transcriptional cross species map of pancreatic islet cells. Molecular Metabolism, 66: 101595, 2022. doi: 10.1016/j.molmet.2022.101595.

X. Wang, D. Choi, and K. Roeder. Constructing local cell-specific networks from single-cell data. Proceedings of the National Academy of Sciences, 118(51):e2113178118, 2021. doi: 10.1073/pnas.2113178118.

H. Wickham. Reshaping data with the reshape package. Journal of Statistical Software, 21(12):1–20, 2007. doi: 10.18637/jss.v021.i12.

H. Wickham. ggplot2: Elegant Graphics for Data Analysis. Springer-Verlag New York, 2016. ISBN 978-3-319-24277-4. URL https://ggplot2.tidyverse.org.

H. Wickham, R. François, L. Henry, K. Müller, and D. Vaughan. dplyr: A Grammar of Data Manipulation, 2023. URL https://CRAN.R-project.org/package=dplyr. R package version 1.1.4.

H. Wickham, D. Vaughan, and M. Girlich. tidyr: Tidy Messy Data, 2024. URL https://CRAN.R-project.org/package=tidyr. R package version 1.3.1.

H. Wickham, T. L. Pedersen, and D. Seidel. scales: Scale Functions for Visualization, 2025. URL https://CRAN.R-project.org/package=scales. R package version 1.4.0.

N. Xiao. ggsci: Scientific Journal and Sci-Fi Themed Color Palettes for ‘ggplot2’, 2024. URL https://CRAN.R-project.org/package=ggsci. R package version 3.2.0.

L. Zappia, B. Phipson, and A. Oshlack. Splatter: simulation of single-cell RNA sequencing data. Genome Biology, 18(1):174, 2017. doi: 10.1186/s13059-017-1305-0.

A. Zeevi and R. Mashal. Beyond correlation: Extreme co-movements between financial assets. SSRN, 2002. doi: 10.2139/ssrn.317122.

Q. Zhang. Classification of RNA-seq data via Gaussian copulas. Stat, 6(1):171–183, 2017. doi: 10.1002/sta4.144.

Q. Zhang and X. Shi. A mixture copula Bayesian network model for multimodal genomic data. Cancer Informatics, 16:1176935117702389, 2017. doi: 10.1177/1176935117702389.

